# The Acheulean niche: Climate and ecology predict handaxe production in Europe

**DOI:** 10.1101/2024.07.19.604259

**Authors:** Michela Leonardi, Stephen J. Lycett, Andrea Manica, Alastair Key

**Affiliations:** Department of Zoology, University of Cambridge, Cambridge, UK; Natural History Museum, London, UK; Department of Anthropology, University at Buffalo (SUNY), Amherst, New York, USA; Department of Archaeology, University of Cambridge, Cambridge, UK

**Keywords:** Lower Paleolithic dispersal, Climate, Paleoenvironment, Ecocultural niche modeling, Prehistoric demography

## Abstract

The Acheulean was the most spatially and temporally vast stone tool industry produced by early humans, persisting for >1.6 million years across three continents. Understanding how behavioral and artifactual variation in these populations relates to climate change and ecology is vital to correctly interpreting the archaeological record, especially given the major climatic and cultural fluctuations during the Pleistocene. Bifacially flaked core tools – which technologically define the tradition – are sometimes regionally and periodically absent within the Acheulean’s boundaries. Biface presence is broadly associated with, but rarely formally tested against, environmental conditions. Here, we investigate biface presence/absence and climate/ecological variation at a continental scale between 726 and 130ka. Using a comprehensive sample of European sites matched to paleoenvironmental reconstructions, we demonstrate clear associations between Acheulean presence and specific environments. Acheulean habitat suitability was most strongly associated with mean winter temperatures above -5/-10°C, and precipitation of the driest month above 15 mm. Two major habitats, warm forest/woodland and shrubland, comprised the European Acheulean niche. Acheulean spatial presence was reconstructed through time: during interglacial and glacial periods it covered most of southwestern Europe, but extended along northern coastal routes during interglacials. Central and Eastern Europe always remained unsuitable. The Acheulean was excluded from areas experiencing both low winter temperatures and low summer precipitation, but reached areas including only one of those two climatic extremes, suggesting seasonal migrations to avoid the harshest seasons. These analyses explain current Acheulean site distributions in Europe and suggest strong links between handaxes, specific ecological conditions, and prehistoric demographic patterns.

## Introduction

Bifacially flaked core technologies define the Acheulean techno-complex ^1^: a lithic cultural phenomenon first observed some 1.8 million years ago (ma) in East Africa ^2^. Subsequently, the Acheulean spread widely across Africa, Europe, and Asia, persisting until the introduction of prepared core technologies around 300 thousand years ago (ka), which occurred in a regionally, temporally and technologically variable manner ^1,3-4^. Classic Acheulean technological practices appear to end in Eurasia at some point after 150-100 ka ^5^. Within these exceedingly long and spatially-broad boundaries, substantial variation is present in the technological features observed at Acheulean archaeological sites. Distinct bifacial tool types and forms existed ^6^, while other flake-based stone technologies are represented in diverse ways ^2,3,7,8^. It is, however, handaxes and cleavers that link archaeological sites with the presence of the Acheulean tradition.

Ironically, the ubiquity of Acheulean biface production across three continents and a range of environmental circumstances, has drawn conspicuous attention to pockets of both time and space within which handaxes appear to not have been produced. From the late Oldowan of Africa ^2^, the Movius line debate in Asia ^9^, through to the ‘Clactonian’ and glacial cycles of northern Europe ^10^, the post ∼1.8 ma absence of bifaces has led to intense debate concerning why these tools were, or were-not, present at Acheulean-period archaeological sites (Figure 1). Explanations for biface presence or absence include discussions on the loss of cultural information, the role of lithic technological innovation, hominin demographic influences, raw material considerations, and variation in utilitarian pressures across differing ecologies ^7-14^.

**Figure 1:**
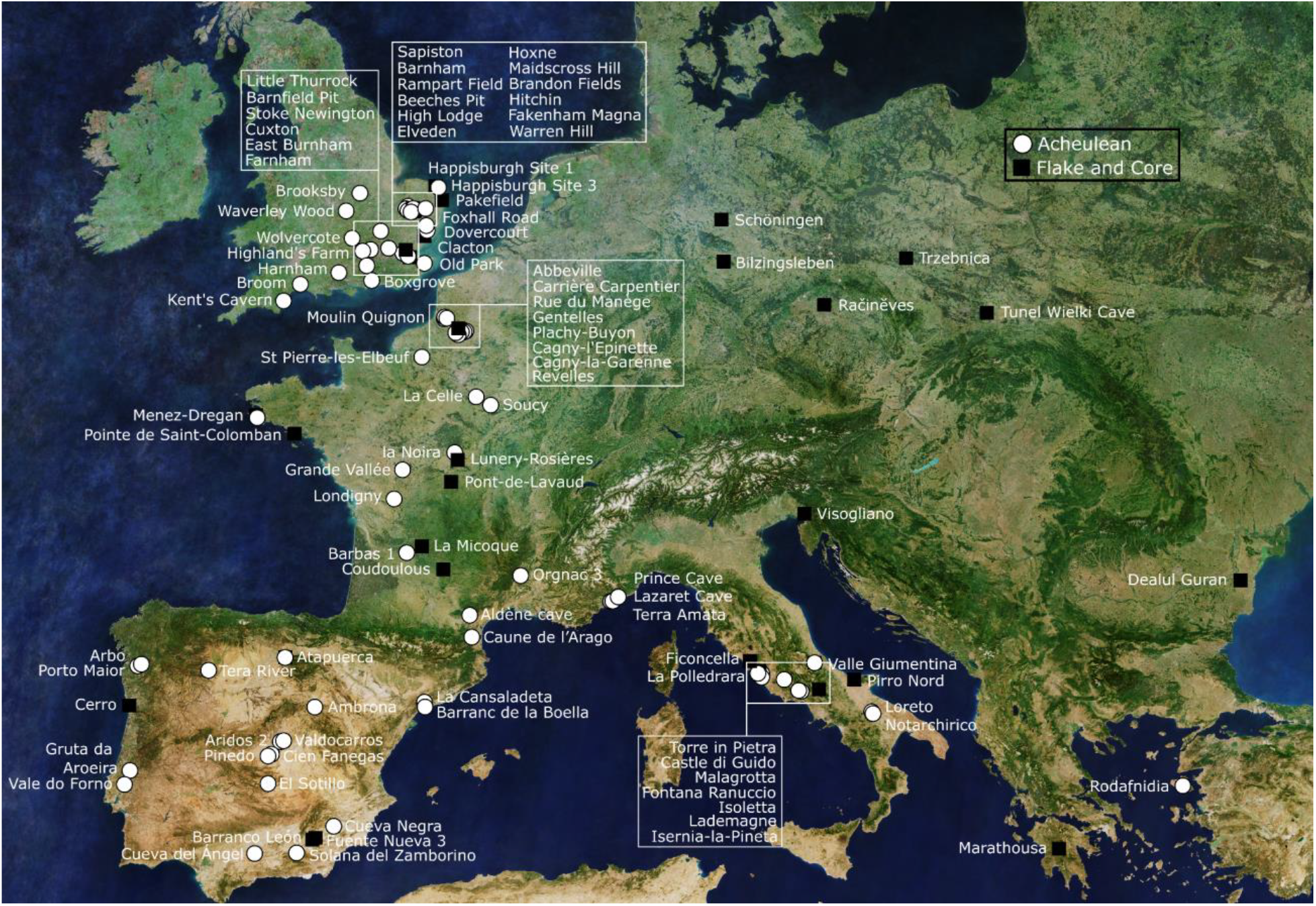
A map detailing all known dated (see methods) Acheulean sites in Europe, along with a proportion of Lower Paleolithic flake-and-core-only sites. Longitudinal and latitudinal data can be viewed in the SI. Note that some sites provide more than one data point due to artifacts being present in multiple stratigraphic layers. In some instances sites contribute both Acheulean and flake-and-core-only data (e.g., Barnfield Pit [UK], Menez-Dregan [France], Atapuerca [Spain]).

Theoretically, we might expect different habitats to have been influential in driving diversity of hominin technological strategies; since bifaces could have given access to important resources in specific environments. Between 1.8 ma and 100 ka, Lower Paleolithic lithic assemblages are observed in a wide range of ecologies — from equatorial rainforests to grasslands, temperate forests, as well as arid, coastal, and high-altitude locations, among others ^10,15,17-19^. Each would have provided distinct challenges, and lithic technologies would have helped hominin populations adapt to these varied niches ^1,7^. If Acheulean biface technologies were required for a population to effectively exploit an ecological niche, then the relevant cultural information would likely have been retained and actuated. If a population’s niche did not necessitate the use of bifaces, then biface-relevant cultural information would have either been lost or not actively practiced ^1,9.^ Climatic and environmental variation could also have indirectly influenced biface presence through their effect on population demography. The suitability of certain habitats to support larger populations, for example, may have been influential, and some ecological conditions – particularly in northern latitudes – may not have been capable of maintaining sizes sufficient to retain complex (i.e., bifacial LCT) cultural information ^10,11,20^. Other demographic factors, such as population bottlenecks occurring during dispersals, could also have influenced biface production ^12^.

Deciphering the impact of ecological variation on the technological behaviors observed at Lower Paleolithic archaeological sites is difficult. Usually, individual site-level inferences are made depending on the artifact types observed and the faunal, floral, and climatic signals present (e.g., ^13,16,18^). At times, ecological information from multiple sites are combined at a local or regional level (e.g., ^21-23^). There are few substantial and structured reviews or data-led studies investigating Lower Paleolithic technological presence or absence at a continental, global or ‘whole industry’ level. We are, then, currently lacking the ability to accurately and more formally link technological variation during the Lower Paleolithic with broad-scale trends in the hominin ecological niche.

In recent years, the reconstruction of paleoclimate information at global/continental scales and across hundreds of thousands, or even millions of years has become possible (e.g. ^24-26^). Thanks to these efforts, researchers can now access diverse bioclimatic variables from across the globe at a millennium-scale, stretching back millions of years. Depending on the reconstruction considered, it is now possible to explore climatic and orography variables, land masses, extent of permanent ice and vegetation/biomes. Here, we use large-scale paleoclimate reconstructions^25^ to investigate the eco-cultural niche of Acheulean bifaces in Europe based on 20 bioclimatic and topography variables, alongside an exhaustive database of dated European Acheulean archaeological sites. We investigate the European Acheulean as it displays the highest resolution data for biface sites in the world. Europe also provides prominent spatially and/or temporally structured instances of biface absence, including during glacial cycles in northern latitudes, in central and eastern European regions, between 800 and 600 ka in Iberia, and a series of MIS 11 flake and core-focused archaeological sites in western regions.

To identify relationships between Acheulean bifaces in Europe and environmental variables, we used eco-cultural niche modeling (ECNM) ^27^. ECNM builds on a method called ecological niche modeling (ENM) ^28-30^ that reconstructs the relationship between species and the climate/environment they inhabit (climatic niche). Through ENM, species presence in one period or area can be used to predict potential distributions when no sighting (occurrence) data are available. ECNM does not focus on biological populations, but cultural ones (i.e., groupings of organisms practicing specific cultures, traditions, or socially-learned behaviors). The assumption underpinning ECNM is that cultural evolutionary mechanisms are impacted by, and have a direct relationship with the environment; and vice versa ^27^.

## Methods

ECNM requires occurrences of interest to be linked with climate records. For prehistoric cultural occurrences – in this case, European Acheulean biface sites – reconstructed paleoclimatic information for each point in space and time is required. We undertook a comprehensive review of Acheulean sites in Europe ^31^, defined as every site on the continent with the confirmed presence of Acheulean bifaces up to the western borders of Russia and Anatolia (Figure 1). Associating each site with paleoclimatic reconstructions requires site-specific spatial and temporal data, while the focus of our investigation necessitated the collection of artifact technological presence and absence information. We therefore excluded archaeological sites from the final dataset if they lacked one or more of these data. Most often, this resulted in exclusions due to a lack of, or lack of reliability concerning, chronological information. Sites or layers defined as ‘Acheulean’ by an article’s authors but also containing evidence of prepared core technologies were excluded. Where dating methods resulted in marine isotope stage (MIS) date assignments, MIS boundaries follow Lisiecki and Raymo ^32^. Every effort was taken for the review to be exhaustive, but we acknowledge a small proportion of sites may have been missed.

Due to the subjectivity of dating reliability determinations, each site was graded between three (highly reliable) and zero (not reliable) on the comprehensiveness of the methods used and whether age assignments are widely accepted. This resulted in two versions of our data; a ‘high reliability’ Acheulean biface site dataset with greater temporal resolution (*n* = 67) and a larger dataset including sites with robust dating methods but lower temporal resolution or acceptance (*n* = 82) (see: supplementary information [SI]). We also collected data from 44 European Lower Paleolithic sites that lack bifaces. This represents a large proportion of the reliably dated flake-and-core-only occurrences in Europe, but our review was not exhaustive. In total, 145 Lower Paleolithic sites are listed in the attached SI dataset, 120 of which display sufficient temporal, spatial and technological data to contribute to our analyses (Figure 1). From each occurrence, we recorded the site name, presence or absence of bifaces, latitude and longitude, upper and lower age thresholds, mean or ‘main’ proposed date (following Key et al. ^6^), dating technique(s), date reliability/acceptance (scaled 0-3), and source citations (see: SI).

We used the R package *pastclim* v. 2.0 ^33^ to extract paleoclimatic and paleoenvironmental reconstructions ^25^ associated with each site’s location and age. The R package *tidysdm* v.0.9.0.9004 ^34^ was then used to perform ECNM based on the aforementioned data. The method requires presence and absence data, but because “real” (i.e., confirmable) absences are not available in the archaeological record, the ‘background’ (spatial and temporal range) of the known data points was randomly sampled to create a set of ‘pseudo-absences’; intersections of spatial and temporal coordinates where the Acheulean is not present ^35^. Here, the background was defined as the whole of Europe, using the above-described spatial limits, between 726 and 130 ka. These dating thresholds represent the upper and lower range limits of the oldest and youngest Acheulean occurrences in our dataset, respectively (Supplementary Information). The two earliest known European Acheulean occurrences, Cueva Negra and Barranc de la Boella (Spain) ^14^, could not be considered as they predate the 800 ka upper limit of the Krapp et al. ^25^ paleoclimatic and paleoenvironmental reconstructions used.

The modeling was performed on two sets of Acheulean data. The first represents Acheulean archaeological sites where published dating evidence was determined to have a reliability score equal to three (n = 65). The second data set included all Acheulean sites with dating reliability greater than zero (i.e., determined to be one, two or three) (n = 80) Each of the subsets was investigated independently. Results obtained from both datasets are in accord. Results based on the latter dataset are presented in Supplementary Information 1 - 4. Seasonal suitability analyses have only been performed on the former set of data in order to be more conservative.

To account for the chronological uncertainty associated with each Acheulean occurrence (represented by associated dating ranges), instead of using the central – often mean – age, a different repetition of the model was run 100 times by sampling from each site’s known temporal range following a normal distribution characterized by the mean ± 2-sigma ^36^ (Supplementary Figure 5). To get enough information on the background, pseudo-absences were added on each time slice by randomly sampling 200 points for each presence and with 70 km as the minimum distance from any presence.

Using *tidysdm*, ecocultural niche modeling was performed separately for each repetition of these differently-dated presences and associated pseudo-absence data points. Each data point was associated with the bioclimatic variables and biome reconstructed for its respective spatial and temporal coordinates ^25^. All 19 annual bioclimatic variables included, available at a spatial resolution of 0.5° every 1000 years, were considered. Elevation is not included in the analyses because when performing ecological modeling through time its effects get confused with temperature ^28^. To account for the effect of topographic variation, rugosity, a measure of altitude variation in a given area, was used instead.

A high level of correlation between bioclimatic variables can bias paleoecological models. The correlation between each pair of variables was calculated, and whenever two showed a value above 0.8, one of them was removed (Supplementary Figure 6). Consequently, the final set of uncorrelated variables used in the models included: mean temperature of the wettest quarter (BIO8), mean temperature of the warmest quarter (BIO10), mean temperature of the coldest quarter (BIO11), precipitation of the wettest month (BIO13), precipitation of the driest month (BIO14), precipitation seasonality (BIO15), precipitation of the warmest Quarter (BIO18), precipitation of the coldest quarter (BIO19), leaf area index (LAI), net primary productivity (NPP), and rugosity.

The models were run independently for each repetition as follows. Spatial block cross-validation was performed using 10 spatial blocks ^37^. The models were tuned with the following algorithms: maxent, generalized linear models (GLM), generalized additive model (GAM), random forest (RF), and generalized boosted models (GBM), exploring 10 different combinations of hyperparameters per model. The fit of each model was evaluated using Boyce Continuous Index (BCI) ^38^, which is specifically designed to be used with presence-only data.

The results for all repetitions were combined by creating a single stacked ensemble ^39^ using the median. Variable importance (how much each variable played a role in shaping the results of each ensemble) was calculated using the DALEX package ^40^.

These final ensembles (one for each subset) were then used to transform the predicted probabilities of occurrence into presence/absence. Different thresholds (0.9, 0.95 and 0.99) were used to map the potential distribution of the Acheulean culture in Europe based on the minimum predicted area encompassing respectively 90%, 95% and 99% of presences. These respectively identify a “core area” (90%), “peripheral area” (95%) and “total area” (99%) suitable for Acheulean populations. These maps were created every 10,000 years (130,000, 140,000 ka, etc.) and for each MIS upper and lower peaks identified in the LR04 stack of benthic 𝛅 ^18^O records ^32^ (Supplementary File 2).

To see the impact of the two most important variables (BIO11 and BIO14) on the model distribution, a scatterplot was created for all time-slices, with each point representing a cell (the spatial unit of the maps, i.e. a “pixel”), differently coloured based on the area they belong to (core, peripheral, total or unsuitable). The plot showed two specific thresholds in BIO11 (below 0 or -10 °C depending on the suitable area considered) and BIO14 (below 15-20 mm depending on the area considered). Areas below both thresholds are considered inhabitable, while areas below only one are considered inhabitable, suggesting seasonal use of these areas. Such seasonally inhabitable regions have been plotted through time similarly to what has been done for the spatial distribution of suitability (Supplementary File 1; Figure 5).

## Results

### The distribution of the European Acheulean

We reconstructed the potential distribution of the European Acheulean through time using an ensemble approach (see Methods for details) and represent the suitable area by considering three different thresholds: a core area (encompassing 90% of the chronologically resampled presences), a peripheral area (95% of presences) and a total area (99% of presences). The modeled Acheulean distribution covers most available landmass in what is now France, the Italian peninsula, mid-to-southern Britain and the Iberian Peninsula (Figure 2). To a lesser extent, the Acheulean was observed in the mid-to-southern Balkans, and coastal regions of Anatolia. The Iberian Peninsula was mostly suitable during glacial periods, but to a lesser extent during interglacials. During interglacials, the tradition’s potential distribution extends north-east along the Atlantic coast. Central and Eastern Europe (e.g. present-day Germany) always remained unsuitable, along with most of Scandinavia. These ranges align with the known distribution of Acheulean sites in Europe (Figure 1).

**Figure 2:**
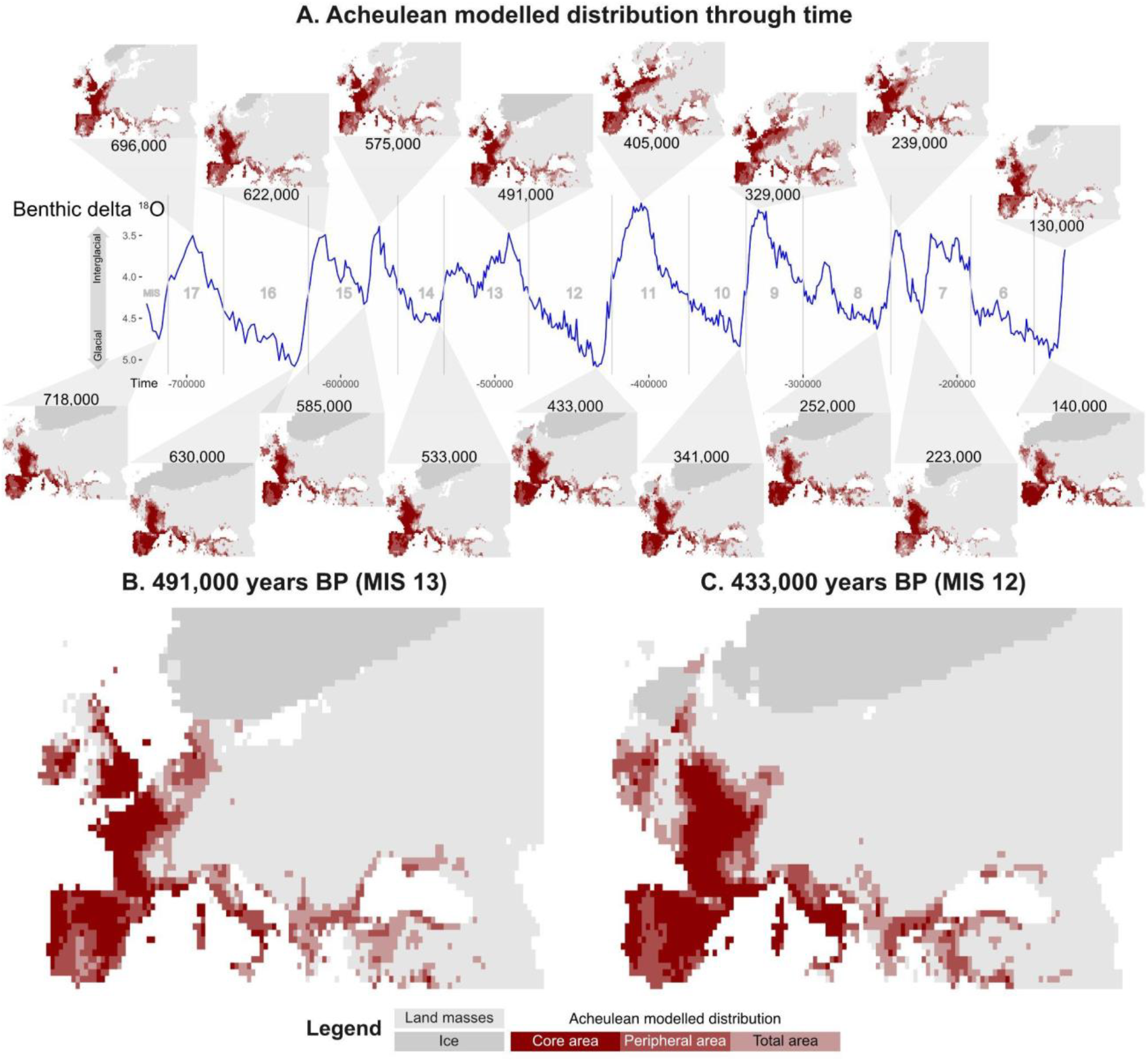
A: The Acheulean’s modeled distribution in Europe for each MIS peak. The blue line represents the benthic delta 18 oxygen stack from Lisieki and Raimo (2005). Note that the y-axis scale is reversed to have warmer (interglacial) periods on the top. B: Detail of the map from 491,000 BP (MIS 13). C: Detail of the map from 433,000 BP (MIS 12). Each map depicts the potential distribution for the core, peripheral and total areas (i.e. covering 90%, 95% and 99% of the presences) respectively, from darker to lighter red.

### Which eco-climatic variables most strongly predict this distribution?

The extent of the suitable range was principally driven by the mean winter temperature (mean temperature of the coldest quarter, BIO11), as demonstrated by the estimate of the variable importance (Figure 3A). A series of additional variables played a less critical, but still substantial, role in the model: three measures of precipitation (driest month [BIO14]; seasonality [BIO15]; and winter [coldest quarter, BIO19]), and one of temperature (mean summer temperature [warmest quarter, BIO10]).

**Figure 3:**
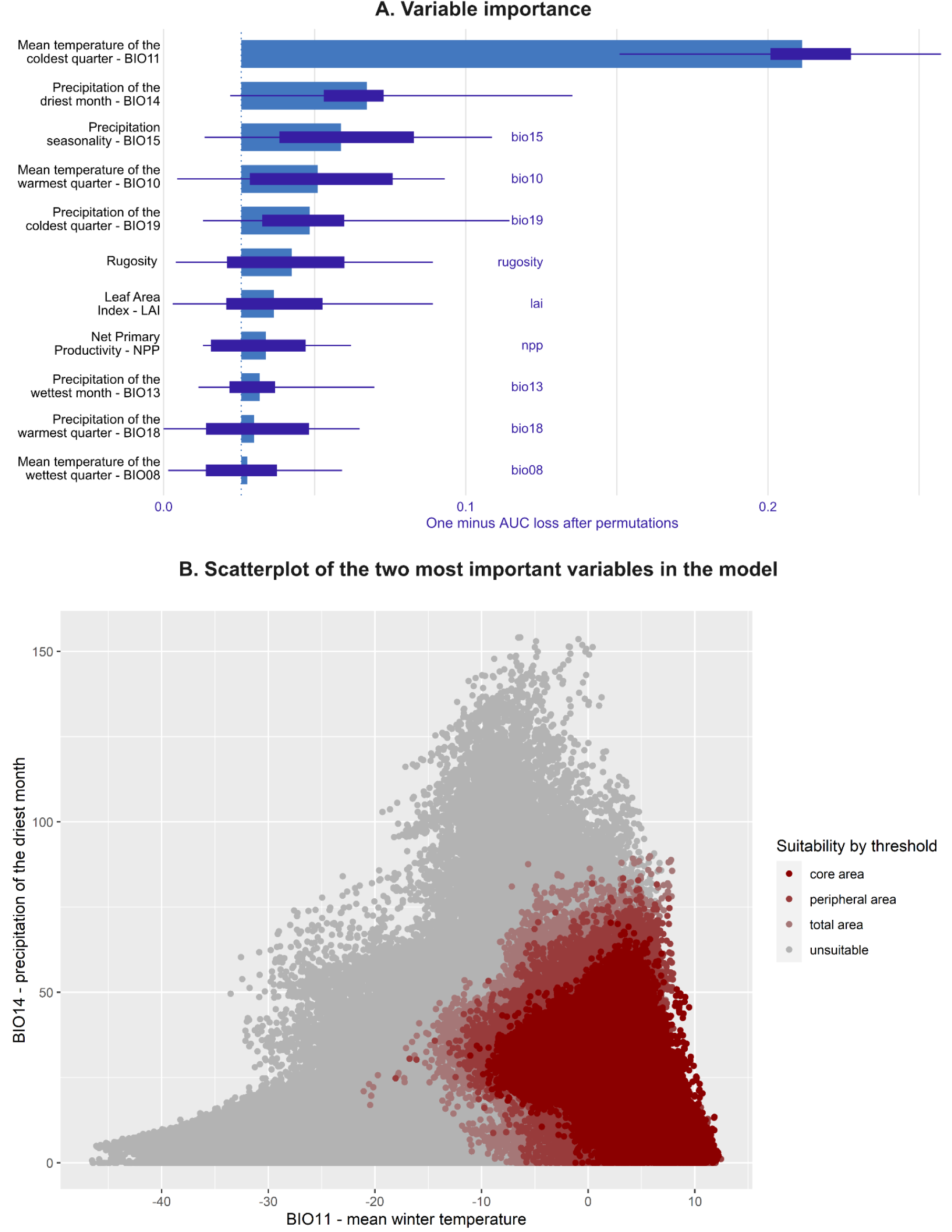
Effect of different variables on the model outcome. A: Variable importance, an estimate of each variable’s contribution to determining the ensemble results. B: Scatterplot of the values of the two most important variables in the model (BIO11 and BIO14), including all time-slices. Each point represents a cell, the colors follow Figure 2 and represent the suitability estimated by the model: unsuitable cells are gray, while from darker to lighter red respectively show suitability based on 90%, 95% and 99% of the presences.

The relationship between the model’s prediction and each variable is depicted in more detail in Supplementary Figure 1. Acheulean suitability (presence) was linked primarily to the mean winter temperature (BIO11) being above -15 °C and precipitation during the driest month (BIO14) being below 80 mm. Suitability was also linked to low precipitation seasonality (BIO15), although, being a coefficient of variation, it should not be considered an absolute value. As high as possible winter precipitation (BIO19) and as low as possible mean summer temperature (BIO10 - especially below 10° C) also increased an area’s suitability for the Acheulean. The interplay among these variables means that, whilst the area suitable for the Acheulean was greater during interglacial than glacial periods, the fluctuations are slightly out of synchrony with the MIS peaks and throughs (which are based on Oxygen18 and capture global temperatures; Supplementary Figures 2 and 3).

### Which environments are suitable for Acheulean populations?

The reconstructed niche of the European Acheulean (core and peripheral areas) (Figure 4), comprises three major habitat types: forest/woodland, shrubland, and tundra. The former covers the vast majority of the Acheulean’s modeled distribution, followed by shrubland in more southern regions of Europe. The Acheulean was present in only a few areas of tundra in Northern Europe – potentially ecotone regions. As tundra covers a negligible part of the suitable area, we do not investigate this environment in more detail.

**Figure 4:**
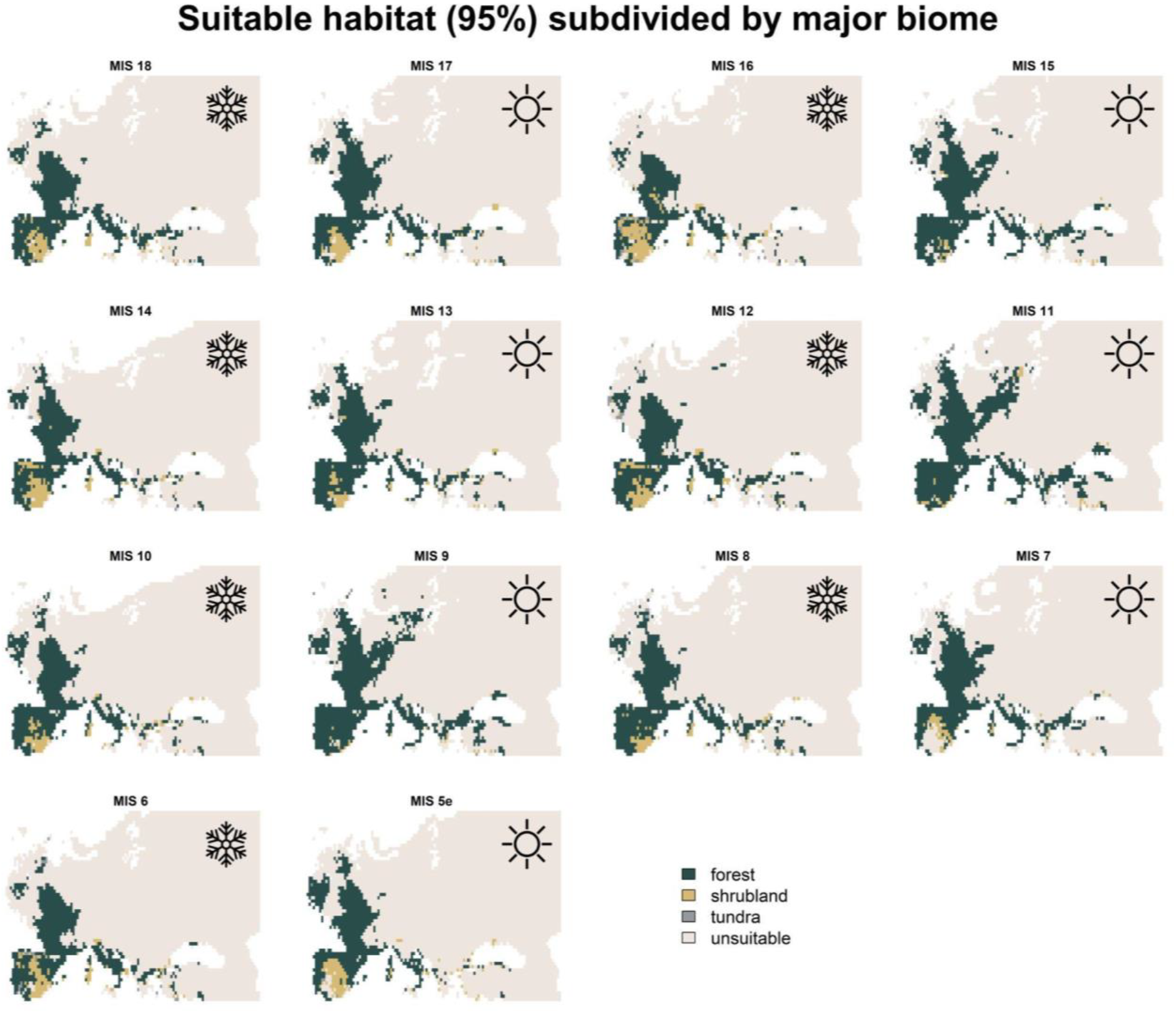
Modeled potential core and peripheral distribution for the Acheulean in Europe subdivided into major habitats: forest (dark green), shrubland (beige), and tundra (gray). Interglacial periods are marked by a sun in the top right corner, while glacial periods by a snowflake.

The most suitable biomes for the Acheulean include areas of temperate/warm forests; including temperate sclerophyll woodland, warm mixed forests and temperate conifer forests, but only rarely temperate deciduous forests (Supplementary Figure 4). The only suitable habitats among colder forests were cool conifer and evergreen taiga forest, and to a lesser extent cool mixed forests and open conifer woodland. While this pattern was consistent in glacial and interglacial periods, during the former, we see slightly higher suitability of colder forests, while warmer forests are slightly more suitable during the latter. Cold mixed forest, deciduous taiga forest and northern tundra were rarely suitable. Only a proportion of available European shrubland is considered suitable, but this biome was more often suitable during interglacial periods (especially temperate xerophytic) (Supplementary Figure 4). If we look at the core area alone, it mostly encompasses temperate sclerophyll woodland and warm mixed forest, with a limited expansion towards neighboring cold mixed forests, suggesting a clear preference for warmer habitats.

### Possible seasonal mobility in Acheulean populations

The most important variables underpinning the model mostly reflect the harshness of winter (mean temperature of the coldest quarter, BIO11) and summer (precipitation of the driest month, BIO 14, limiting water availability). Because each of them is informative for a different, extreme, season, they can help us understand what climatic challenges Acheulean populations could withstand at different times of the year. Looking at the values for these two variables for cells that are predicted as suitable (Figure 3B, red points) it is possible to see that Acheulean groups could live in areas with cold winters (y-axis, BIO11, below 0 or -10 °C, depending on the threshold considered). They could also withstand extremely dry summers (y-axis, BIO14, below 15-20 mm/month). But the plot shows how cells where both variables fall below the above-mentioned thresholds are considered unsuitable. In other words, areas with a cold winter but a relatively wet summer were predicted to be suitable, as were areas with a mild winter and a dry summer. This suggests the possibility that these areas were used only during the favorable part of the year, thus implying seasonal mobility.

Figure 5 shows the spatial distribution through time of such seasonally habitable regions, differently shaded based on them being part of the core, peripheral or total area. Areas with very cold winters and rainy driest months are observed in north-central Europe, while those with warmer winters but low precipitation in the driest month can be found in most of the Iberian Peninsula. We investigated the suitability associated with five important Acheulean sites in detail: two from northern Europe (Maidscross Hill and Beeches Pit); one from Central Europe (La Noira lower), and two from Southern Europe (Vale do Forno Upper sands, and Solana del Zamborino) (Figure 5D). While La Noira is located in an area favorable to year-round occupation during both glacial and interglacial phases, Maidscross Hill and Beeches Pit may have been unfavorable for winter occupation during portions of glacial periods, pointing to the potential of seasonal migrations during the coldest phases. Southern sites are found in regions for which our model considers the summer is too dry to allow Acheulean occupation, more strongly supporting seasonal migrations in this instance.

**Figure 5:**
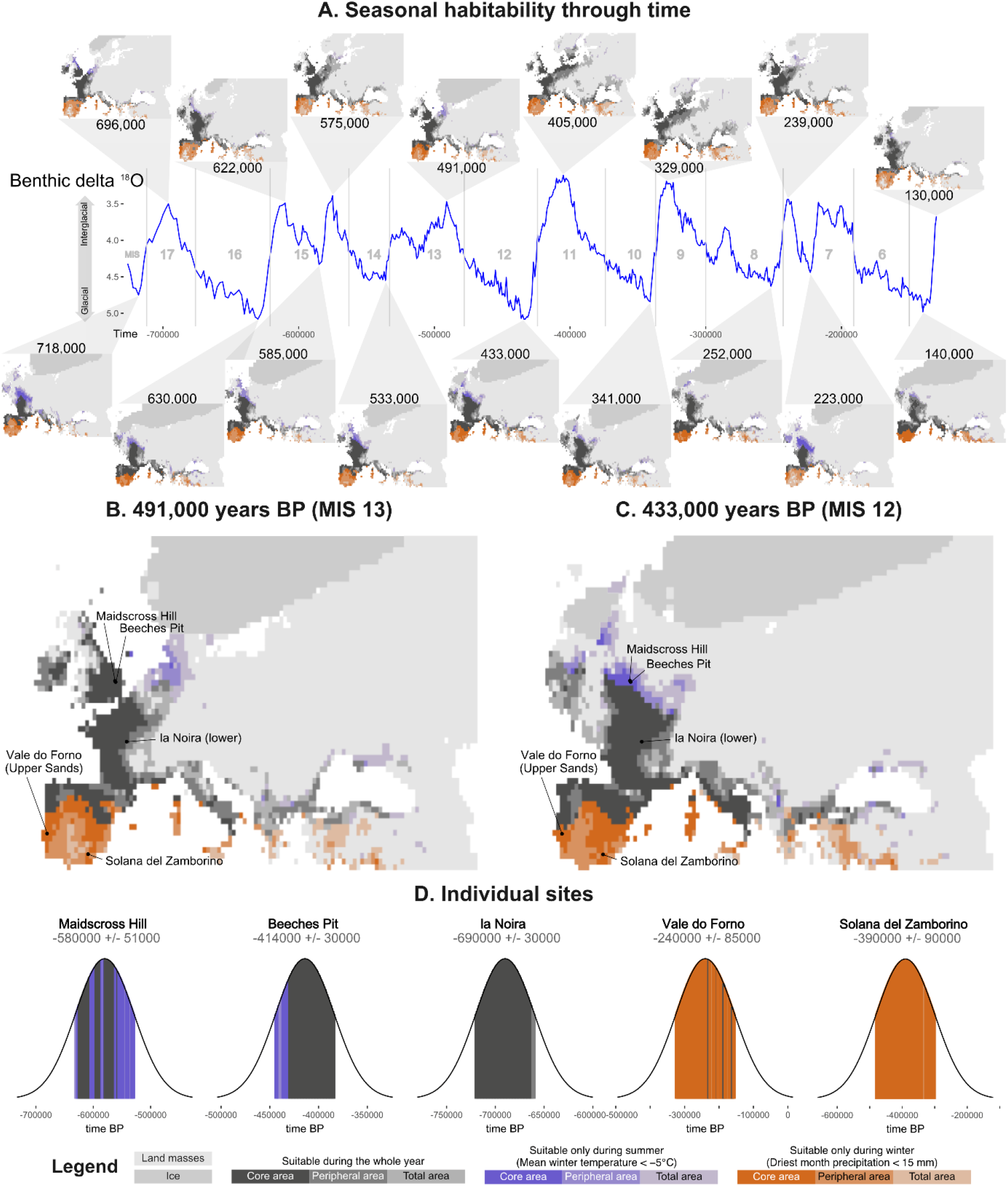
A: Modeled distribution of areas suitable for Acheulean populations on a seasonal basis. Southern orange areas indicate locations that become too arid during the summer for hominins, while blue regions were potentially too cold for Acheulean populations during the winter. Note that the strongest high-latitude signal for these potential seasonal migrations is during glacial periods. B and C: Modeled distribution during the MIS 13 interglacial and MIS 12 glacial periods, noting the four potential migration sites (Maidscross Hill [unsuitable in winter], Beeches Pit [unsuitable in winter], Vale do Forno [unsuitable in summer] and Solana del Zamborino [unsuitable in summer]), alongside la Noira as a representative site for year-round occupation.

## Discussion

Climate and ecology help explain diverse aspects of human behavior, in the present day and the deep past. Temporal and spatial variation in the Acheulean stone tool phenomena, the single longest cultural tradition to have been produced by the hominin lineage ^1,6^, has long been linked to Pleistocene climatic and ecological change (e.g., ^41^). Rarely have these hypotheses been formally tested, and never against continental-scale paleoenvironmental reconstructions. Here, we demonstrate strong links between Acheulean site distributions in Europe and specific paleoclimatic variables.

The distribution for Acheulean culture in Europe, modeled using paleoclimatic data, closely matches its known, archaeologically evidenced distribution (Figure 1), demonstrating climate and ecology play an important role in determining its presence during the Pleistocene. The ecocultural niche of the European Acheulean is overwhelmingly associated with warm-to-temperate forest (incl. woodland) biomes, and shrubland in more southern, Mediterranean regions. Driving this association are mean winter temperatures, with Acheulean populations displaying lower thresholds of -5 to -10 °C, alongside sensitivity to precipitation during the driest month, having a lower threshold of 20-15 mm/month. Extreme climate regions, where one of these variables is below its threshold, are predicted as habitable when the other variable has higher values. Low seasonal variation in precipitation, high precipitation during winters, and relatively cool mean summer temperatures also increase the likelihood of Acheulean site presence.

These climatic drivers explain why the Acheulean’s modeled distribution is represented by a ‘focused area’ covering most of Iberia, present-day France, Britain, the Italian peninsula and coastal areas in the Balkans and Anatolia. The northern extension of the Acheulean’s modeled range along the warm Atlantic coast emphasizes the role of mean winter temperatures, with Acheulean distributions pushed to southern Scandinavia during interglacial periods (Figure 2). High levels of precipitation along Atlantic coastal regions would have strengthened this distribution trend, making colder mean winter temperatures tolerable (Figures 2 and 5). Potentially, the expansion of the dry Mammoth steppe in the Late Acheulean explains why the latest known European sites are all in southern France and Iberia ^31^. Reduced temperature annual range resulted in increased habitable areas, potentially indicating Acheulean populations to better cope with stable climates relative to those that are more seasonally variable.

Middle Pleistocene eastern and northern Europe, where there is a stark absence of Acheulean sites (Figure 1), is characterized by cold mixed forest, cool conifer forest, and tundra during colder glacial stages, with temperate conifer and cool mixed forests increasing during interglacial periods (Supplementary Figure 4). These biomes appear mostly unsuitable for Acheulean populations within their European range ^20,23^, yet several appear habitable when they occur in more westerly and southerly regions, often during glacial periods. An unequivocal link between these habitat types and the presence or absence of Acheulean cultural information is, therefore, unsupported. In other words, there are few biomes that appear completely unsuitable for the retention and/or application of Acheulean cultural information, even if only for spatially or temporally limited periods. Hence, Acheulean technologies appear to have been useful to hominins in almost any habitat ^15^, although Acheulean populations were clearly warm forest and woodland specialists.

From a utilitarian perspective ^7^ it is difficult to see how these specific ecologies would restrict Acheulean cultural information to the ranges in Figure 2. Potentially, the medium-to-large fauna that bifaces are associated with butchering represented a higher percentage of the biomass in southern and western regions, but such fauna are not rare or absent in the north or east ^23,42^. Some bifaces make excellent digging tools, but plant tubers, eusocial insects and burrowing animals are present in cold, warm, dry and wet environments. Permafrost would have potentially impacted digging behaviors, but its range does not always mirror the Acheulean’s modeled boundaries ^43^. Finally, bifaces effectively modify plant materials ^7^, but Middle Pleistocene organic tools are occasionally found in Eastern regions and suitable source materials would have been present in almost all biomes ^44^. Even if differences between biomes did reduce the frequency of their use, bifaces would still be present in the Paleolithic record at low frequencies. At present, evidence suggests handaxes were a flexible technological strategy, suited to a wide range of ecologies, wherever the recurring need for a heavy-duty cutting implement presented itself ^7^. Indeed, the ubiquity of handaxe production across swathes of time and space might be taken to mean that handaxes were produced whenever Middle Pleistocene hominins had the cultural knowledge to do so.

What instead appears a more likely driver of Acheulean spatial presence was the impact of temperature and precipitation on hominin demography, rather than a direct utilitarian impact of ecology on handaxe production. When populations were relatively smaller, there was a greater likelihood of cultural information loss due to a reduced number of skilled individuals to learn from and a heightened potential for cultural drift to take effect ^12,45^, especially in a relatively complex activity such as biface production ^46^. Sites containing only core-and-flake assemblages during the Acheulean period — including western and southern regions — may, therefore, be the product of relatively smaller hominin populations restricted by ecology (e.g., ^22,23^).

Mean winter temperature has long been argued to be a limiting factor on European Acheulean population northern distributions and densities ^21,22,43^, but it is only now that a lower threshold of -5 to -10 °C is evidenced. This threshold was potentially mediated by technological adaptations (e.g., use of fire, clothing, and so forth ^42,47^) and the limitations of Acheulean populations to deploy such strategies effectively. Importantly, because BIO11 is a mean over an entire season, it tells us little about the number of days below the threshold that hominins could have survived. Notably, and supporting past work ^19,20,31^ but in opposition to more established interpretations ^10,21^, western portions of northern Europe appear suitable for hominin habitation during glacial periods (Figure 2). Low summer precipitation has similarly been hypothesized to limit Pleistocene hominin population distributions ^16,17^, and the ethnographic literature also indicates this will restrict human population ranges ^48^. Reliable access to water was essential for hominins ^47^, with summer precipitation potentially having a year-round impact on groundwater levels dependent on local geology. This could help explain the unsuitability of southern Iberia during the peaks of interglacial periods (Figure 2).

As colder (−15 °C) areas are only habitable with greater precipitation during the summer, the joint impact of temperature and precipitation on hominins could be explained in two ways. On the one hand, hominins could cope with either cold winters or dry summers, but not when both are experienced in the same region. Alternatively, and arguably more likely, these areas may only have been used by Acheulean hominins during favorable periods, with populations relocating to nearby suitable regions when temperature or precipitation conditions became too harsh. This latter scenario suggests Acheulean hominins in the northern and southern extremes of their European range may have undertook seasonal migrations to avoid low winter temperatures or excessively dry summers, respectively.

From both ethnography and archaeology, we know that seasonal migration is an important strategy in *Homo sapiens*, especially hunter-foragers ^48^. However, in the case of Middle Pleistocene hominins, the issue of seasonal migration is nearly impossible to infer directly from the archaeological record, since chronology of occupation is too imprecisely measured. Even evidence from a site or sites demonstrating the exploitation of seasonally available resources, does not necessarily — by itself — allow for indisputable inference of strategic seasonal migration and pre-planned avoidance of specific biomes at certain times of the year. This is important since, within the human lineage at least, evidence of relatively long-distance seasonal migration would imply “planning depth”, or the cognitive ability to strategically plan activities in anticipation of future events and in advance of immediate needs ^49^.

Our analysis provides strong evidence for migration for southern-latitudes In Iberia, the north of the peninsula was always habitable and, at times, suitability expanded towards its center. Southern Iberia was, however, only seasonally suitable across the whole analyzed period, with summers having been too arid for Acheulean hominin occupation (see seasonal suitability of Iberian sites in figure 5D). This can be interpreted as strong evidence of seasonal migration in this region, up to the order of a few hundred kilometers (Figure 5. e.g. Solana del Zamborino). The potential for Acheulean populations to have undertaken seasonal (summer) migrations to exploit northern European resources has been proposed before ^21,42^, but precipitation-driven seasonal migrations to and from southern and central Iberia are rarely discussed. On the other hand, migrations at higher latitudes are less supported. Figure 5 shows that in Northern Europe, areas with a high number of Acheulean sites (e.g., Britain) appear suitable for the whole year during interglacials, and mostly only suitable during the warmest part of the year in glacial periods (Figure 5,) As most northern sites are dated to interglacial periods (Supplementary Data), caution is needed when inferring high-latitude seasonal migrations. Our models suggest the possibility of such migration due to the fact that, once dating uncertainty is accounted for, some time-slices within the possible occupancy range would have only been possible for migratory populations (clearly seen in Maidcross Hill in Figure 5); it is however also possible that Acheulean populations deserted those locations during the occasional, unfavourable periods.

The implications of identifying seasonal migration in early human populations is substantial. In *H. sapiens*, it is known to be a complex behavioral strategy demanding detailed knowledge of landscapes and resources, acute perception of seasonal ecological change, and complex group cooperation ^48^. The distances inferred are also among the farthest evidence for the movement of Acheulean individuals to date. For sites strongly associated with migratory zones, such as Vale do Forno and Solana del Zamborino in Iberia (Figure 5D), artifacts may have been transported substantial distances, potentially explaining proportions of curated vs. expedient tools relative to sites in more seasonally stable areas (e.g., la Noira [lower]), or the presence of unexpected raw materials. For others, such as Beeches Pit (UK), which displays a number of time slices within its range associated with potential migratory zones (Figure 5D), important behavioral evidence, such as the site’s early evidence for controlled fire use, could be linked to culturally-mediated adaptations to challenging environments.

Climatically and ecologically mediated barriers, bottlenecks or stimuli to the flow of cultural information during the Acheulean would have created potential for distinct cultural representations to have developed ^9,10,11,12,46^. The present data indicate multiple potential dispersal bottlenecks along the Mediterranean coast, including between southern France and the Italian peninsula, where only a thin strip of land between the Mediterranean and the Alps appears suitable (Figure 2). This would have severely limited the flow of cultural and genetic information, helping to explain why the Acheulean may have been present in the Italian peninsula during a sustained period of absence in Iberia between c. 800 and 500 ka ^14,31^, and why distinct phenotypes such as *H. antecessor* are only present in western European regions. At a continental level, Acheulean populations may have experienced highly limited dispersal potential between the Levant and Europe (and *vice versa*), with only thin coastal routes appearing viable. Cultural isolation facilitated by rare connection with the Levant may explain the familiar yet distinct technological and morphological attributes of western European bifaces ^8,50^, and why northern Africa could have been a similarly viable route – if also restrictive – for the dispersal of Acheulean culture ^8,15,50^.

Whether the pertinent climatic and ecological conditions identified here, or entirely different climatic and ecological variables, had similar effects upon biface production and the ranges of Acheulean hominins in other continents, such Africa and Asia, awaits future study.

## Acknowledgments

ML and AM were funded by the Leverhulme research grant RPG-2020-317.

## References

1. Kuhn, S.L. 2020. The Evolution of Paleolithic Technologies. Routledge, New York

2. de la Torre, I. 2016. The origins of the Acheulean: past and present perspectives on a major transition in human evolution. Philosophical Transactions of the Royal Society B 371: 20150245

3. Gowlett, J.A.J. 2015. Variability in an early hominin percussive tradition: the Acheulean versus cultural variation in modern chimpanzee artefacts. Philosophical Transactions of the Royal Society B 370: 20140358

4. Moncel, M.-H., Ashton, N., Lamotte, A., Tuffreau, A., Cliquet, D. and Despriée, J. 2015. The early Acheulian of north-western Europe. Journal of Anthropological Archaeology 40: 302–331

5. Key, A.J.M., Jarić, I. and Roberts, D.L. 2021. Modelling the end of the Acheulean at global and continental levels suggests widespread persistence into the Middle Palaeolithic. Humanities and Social Sciences Communications 8: 55

6. Lycett, S.J. and Gowlett, J.A.J. 2008. On questions surrounding the Acheulean ‘tradition’. World Archaeology 40 (3): 295–315

7. Key, A.J.M. and Lycett, S.J. 2017. Form and function in the Lower Palaeolithic: history, progress, and continued relevance. Journal of of Anthropological Sciences 95: 67–108

8. Sharon, G. and Barsky, D. 2016. The emergence of the Acheulian in Europe - a look from the east. Quaternary International 411 (Part b): 25–33

9. Schick, K.D., 1994. The Movius line reconsidered. In: Corruccini, R.S., Ciochon, R.L. (Eds.), Integrative Paths to the Past. Prentice Hall, Englewood Cliffs, NJ, pp. 569–596.

10. Ashton, N. and Davis, R. 2021. Cultural mosaics, social structure, and identity: The Acheulean threshold in Europe. Journal of Human Evolution 156: 103011

11. Mithen, S. 1994. Technology and society during the Middle Pleistocene: hominid group size, social learning and industrial variability, Cambridge Archaeological Journal 4: 3–32

12. Lycett, S.J. and Norton, C.J. 2010. A demographic model for Palaeolithic technological evolution: the case of East Asia and the Movius Line. Quaternary International, 211(1-2): 55–65.

13. Rodríguez-Gómez, G., Mateos, A., Martin-Gonzalez, J.A., Blasco, R., Rosell, J. and Rodríguez, J. 2014. Discontinuity of human presence at Atapuerca during the Early Middle Plesitocene: a matter of ecological competition? PLOS One, 9 (7): e101938

14. Ollé, A., Lombao, D., Asryan, L., García-Medrano, P., Arroyo, A., Fernandez-Marchena, J.L., Yesilova, G.C., Caceres, I., Huguet, R., Lopez-Polin, L., Pineda, A., Garcia-Tabernero, A., Fidalgo, D., Rosas, A., Saladie, P. and Vallverdu, J. 2023. The earliest European Acheulean: new insights into the large shaped tools from the late Early Pleistocene site of Barranc de la Boella (Tarragona, Spain). Frontiers in Earth Sciences, 11: 1188663

15. Candy, I., Schreve, D. and White, T.S. 2015. MIS 13-12 in Britain and the North Atlantic: understanding the palaeoclimatic context of the earliest Acheulean. Journal of Quaternary Science 30 (7): 593–609

16. Ecker, M., Brink, J.S., Rossouw, L., Chazan, M., Horwitz, L.K. and Lee-Thorp, J.A. 2018. The palaeoecological context of the Oldowan–Acheulean in southern Africa. Nature Ecology and Evolution 2: 1080–1086

17. Blain, H.-A., Fagoaga, A., Ruiz-Sanchez, F.J., Garcia-Medrano, P., Olle, A. and Jimenez-Arenas, J.M. 2021. Coping with arid environments: A critical threshold for human expansion in Europe at the Marine Isotope Stage 12/11 transition? The case of the Iberian Peninsula. Journal of Human Evolution 153: 102950

18. Lupien, R.L., Russell, J.M., Subramanian, A., Kinyanjui, R., Beverly, E.J., Uno, K.T., de Menocal, P., Domain, R. and Potts, R. 2021. Eastern African environmental variation and its role in the evolution and cultural change of Homo over the last 1 million years. Journal of Human Evolution 157: 103028

19. Moncel, M.-H., Antoine, P., Herisson, D., Locht, J.-L., Hurel, A. and Bahain, J.-J. 2022. Were hominins specifically adapted to North-Western European territories between 700 and 600 ka? New insights into the Acheulean site of Moulin Quignon (France, Somme Valley). Frontiers in Earth Science 10, doi:10.3389/feart.2022.882110

20. Rodríguez, J., Willmes, C., Sommer, C. and Mateos, A. 2022. Sustainable human population density in Western Europe between 560.000 and 360.000 years ago. Scientific Reports 12: 6907

21. MacDonald, K., Martinon-Torres, M., Dennell, R.W. and Bermúdez de Castro, J.M. 2012. Discontinuity in the record for hominin occupation in south-western Europe: Implications for occupation of the middle latitudes of Europe. Quaternary International 271: 84–97

22. Mosquera, M., Ollé, A., Saladié, P., Cáceres, I., Huguet, R., Rosas, A., Villalaín, J., Carrancho, A., Bourles, D., Brancher, R., Pineda, A. and Vallverdu, J. 2016. The early Acheulean technology of Barranc de la Boelle (Catalonia, Spain). Quaternary International 393: 95–111

23. Szymanek, M. and Julien, M.-A. 2018. Early and Middle Pleistocene climate-environment conditions in Central Europe and the hominin settlement record. Quaternary Science Reviews 198: 56–75

24. Beyer, R. M., Krapp, M., and Manica, A. 2020. High-resolution terrestrial climate, bioclimate and vegetation for the last 120,000 years. Scientific Data 7: 236

25. Krapp, M., Beyer, R.M., Edmundson, S.L., Valdes, P.J. and Manica, A. 2021. A statistics-based reconstruction of high-resolution global terrestrial climate for the last 800,000 years. Scientific Data 8: 228

26. Barreto, E., Holden, P.B., Edwards, N.R., Rangel, T.F. 2023. PALEO-PGEM-Series: A spatial time series of the global climate over the last 5 million years (Plio-Pleistocene). -Global Ecology and Biogeography 32: 1034–1045

27. Banks, W.E., Wohlfarth, B., Sánchez-Goñi, M.-F., Marean, C.W., Crucifix, M., Montet-White, A., Gilliam, J.C., Anderson, D.G., Peterson, A.T., Olszewski, D.I., West, D., Krishtalka, L., Dibble, H.L., d’Errico, F. and Vanhaeran, M. 2006. Eco-cultural niche modeling: new tools for reconstructing the geography and ecology of past human populations. PaleoAnthropology 2006: 68–83

28. Guisan, A., Thuiller, W., and Zimmermann, N. 2017. Habitat Suitability and Distribution Models: With Applications in R (Ecology, Biodiversity and Conservation). Cambridge University Press, Cambridge

29. Banks, W.E., d’Errico, F., Peterson, A.T., Vanhaeren, M., Kageyama, M., Sepulchre, P., Ramstein, G., Jost, A. and Lunt, D. 2008. Human ecological niches and ranges during the LGM in Europe derived from an application of eco-cultural niche modeling. Journal of Archaeological Science 35(2): 481–491

30. d’Errico, F., Banks, W.E., Warren, D.L., Sgubin, G., van Kiekerk, K., Henshilwood, C., Daniau, A.-L., and Sánchez-Goñi, M.-F. 2017. Identifying early modern human ecological niche expansions and associated cultural dynamics in the South African Middle Stone Age. PNAS 114(30): 7869–7876

31. Key, A. 2024. Regional extinction(s) but continental persistence in European Acheulean culture. Cambridge Prisms: Extinction, doi: 10.1017/ext.2024.13

32. Lisiecki, L.E. and Raymo, M.E. 2005. A Pliocene-Pleistocene stack of 57 globally distributed benthic δ18O records. Paleoceanography and Paleoclimatology 20(1): 1-17

33. Leonardi, M., Hallett, E.Y., Beyer, R., Krapp, M. and Manica, A. 2023. Pastclim 1.2: an R package to easily access and use paleoclimatic reconstructions. Ecography 3: e06481

34. Leonardi, M., Colucci, M., Pozzi, A.V., Scerri, E.M.L., and Manica, A. 2024. tidysdm: leveraging the flexibility of tidymodels for species distribution modelling in R. bioRxiv 2023.07.24.550358

35. Barbet-Massin, M., Jiguet, F., Albert, C.H. and Thuiller, W. 2012. Selecting pseudo-absences for species distribution models: how, where and how many? Methods in Ecology and Evolution 3(2): 327–338

36. Will, M., Krapp, M., Stock, J. T. and Manica, A. 2021. Different environmental variables predict body and brain size evolution in Homo. Nature Communications 12: 4116

37. Roberts, D.R., Bahn, V., Ciuti, S., Boyce, M.S., Elith, J., Guillera-Arroita, G., Hauenstein, S., Lahoz-Monfort, J.J., Schroder, B., Thuiller, W., Warton, D.I., Wintle, B.A., Hartig, F. and Dormann, C.F. 2017. Cross-validation strategies for data with temporal, spatial, hierarchical, or phylogenetic structure. Ecography, 40 (8): 913–929.

38. Boyce, M.S., Vernier, P.R., Nielsen, S.E. and Schmiegelow, F.K.A. 2002. Evaluating resource selection functions. Ecological Modelling, 157: 281–300

39. Araújo, M. B. and New, M. 2007. Ensemble forecasting of species distributions. Trends in Ecology and Evolution, 22: 42–47

40. Biecek, P. 2018. DALEX: Explainers for Complex Predictive Models in R. Journal of Machine Learning Research, 19(84): 1-5

41. Isaac, G. 1984. The archaeology of human origins: studies of the Lower Pleistocene in East Africa 1971-1981. In: Wendorf, F.and Close, A. (Eds.) Advances in Old World Archaeology. Academic Press, New York. pp. 1–87

42. Hosfield, R. 2016. Walking in a winter wonderland? Strategies for Early and Middle Pleistocene survival in midlatitude Europe. Current Anthropology 57 (5): 653–682

43. Andrieux, E., Bertran, P. and Saito, K. 2016. Spatial analysis of the French Plesitocene permafrost by a GIS database. Permafrost and Periglacial Processes 27 (1): 17–30

44. Popescu, S.-M., Biltekin, D., Winter, H., Suc, J.-P., Melinte-Dobrinescu, M.C., Klotz, S., Rabineau, M., Combourieu-Nebout, N., Clauzon, G., and Deaconu, F. 2010. Pliocene and Lower Pleistocene vegetation and climate changes at the European scale: Long pollen records and climatostratigraphy. Quaternary International 219 (1-2): 152–167

45. Powell, A., Shennan, S. and Thomas, M.G. 2009. Late Pleistocene demography and the appearance of modern human behavior. Science 324 (5932): 1298–1301

46. Lycett, S. J., Schillinger, K., Eren, M. I., von Cramon-Taubadel, N., & Mesoudi, A. 2016. Factors affecting Acheulean handaxe variation: Experimental insights, microevolutionary processes, and macroevolutionary outcomes. Quaternary International, 411: 386–401.

47. Shea, J.J. 2023. The Unstoppable Human Species. Cambridge University Press, Cambridge

48. Kelly, 2013. The Lifeways of Hunter-Gatherers: The Foraging Spectrum. Cambridge University Press, Cambridge

49. Roebroeks, W., Kolen, J., & Rensink, E. (1988). Planning depth, anticipation and the organization of Middle Palaeolithic technology: the archaic natives meet Eve’s descendants. Helinium 28 (1): 17–34.

50. Moncel, M.H., Arzarello, M., Boeda, E., Bonilauri, S., Chevrier, B., Gaillard, C., Forestier, H., Yinghua, L., Semah, F. and Zeitoun, V. 2018. Assemblages with bifacial tools in Eurasia (third part). Considerations on the bifacial phenomenon throughout Eurasia. Comptes Rendus Palevol 17 (1-2): 77–97

