## Supplementary figures for "The Acheulean niche: Climate and ecology predict handaxe production in Europe"

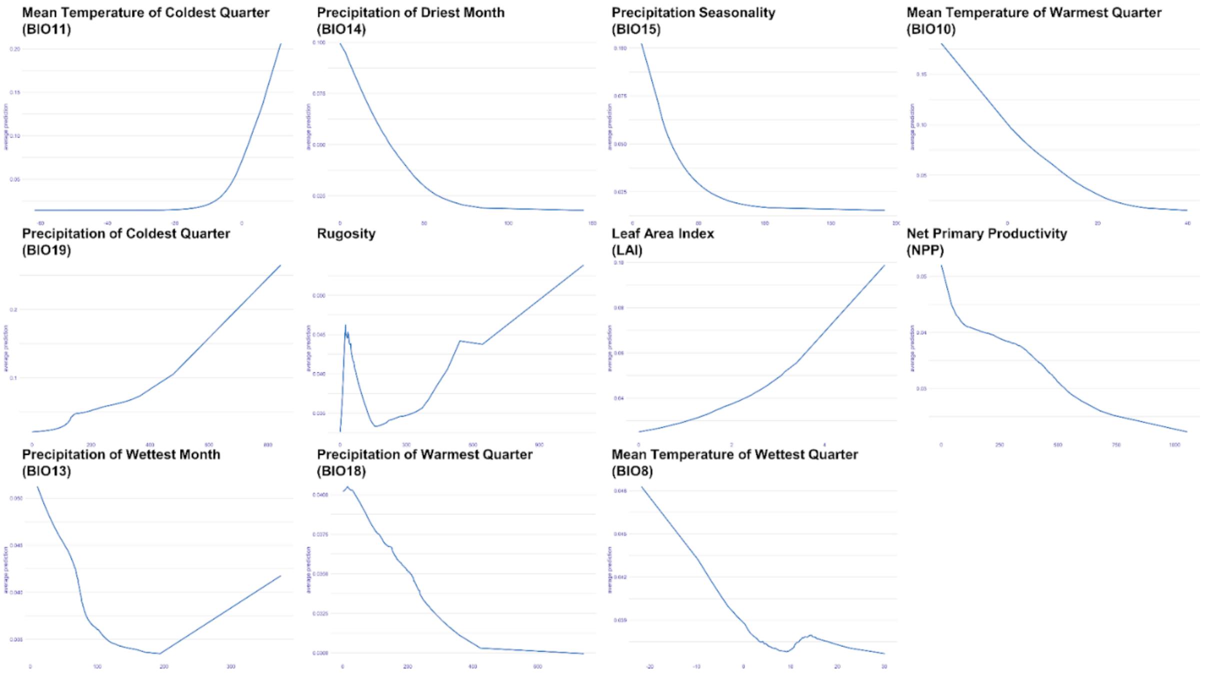


**Supplementary Figure 1**: Partial dependency plots, showing how the expected value of the ensemble prediction (y-axis) behaves as a function of the values of each bioclimatic variable (x-axis). The variables are arranged from the most (top left) to the least (bottom right) important in the model (same as Figure 3).

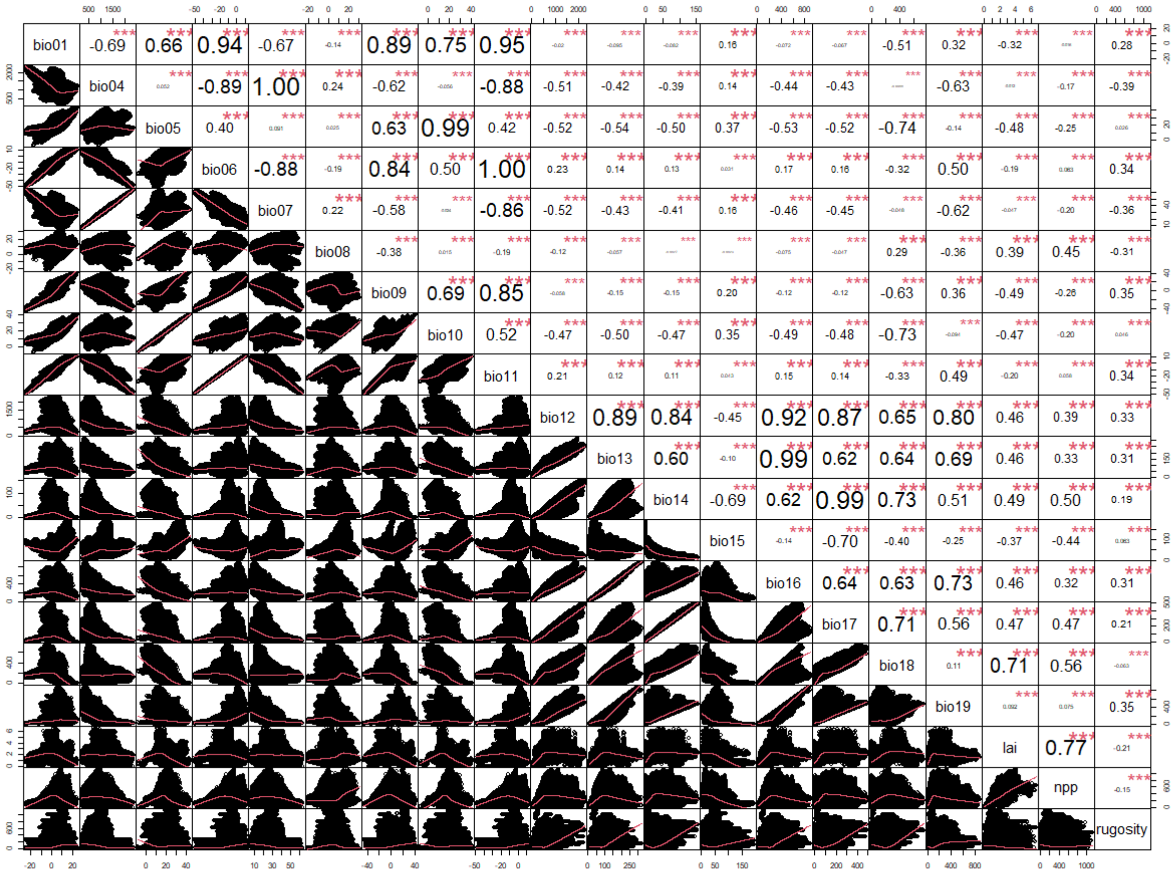

**Supplementary Figure 2**: Cross-correlation of all bioclimatic variables available, across the whole area and period analysed (background).

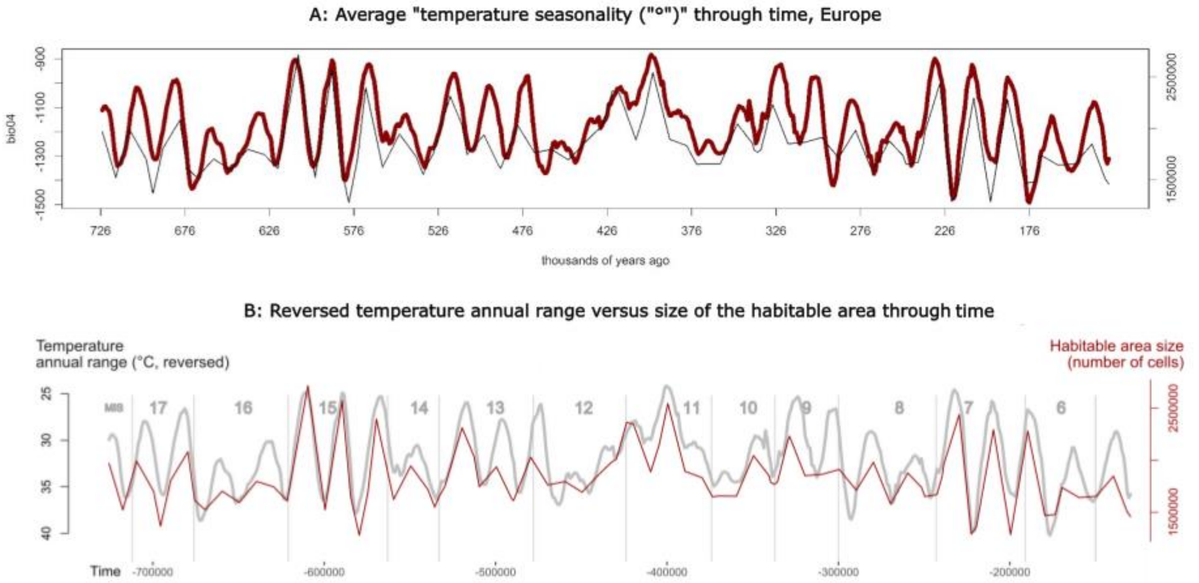


**Supplementary Figure 3**: A: Inverted averaged temperature seasonality (BIO04, expressed as minus standard deviation ×100) over the whole of Europe through time (thick red line, left y-axis), and the number of cells in the core and peripheral areas through time (thin black line, right y-axis). B: Temperature annual range (BIO04) averaged over the whole of Europe through time (thick gray line, left y-axis, reversed), and number of habitable cells in the core and peripheral areas through time (thin red line, right y-axis).

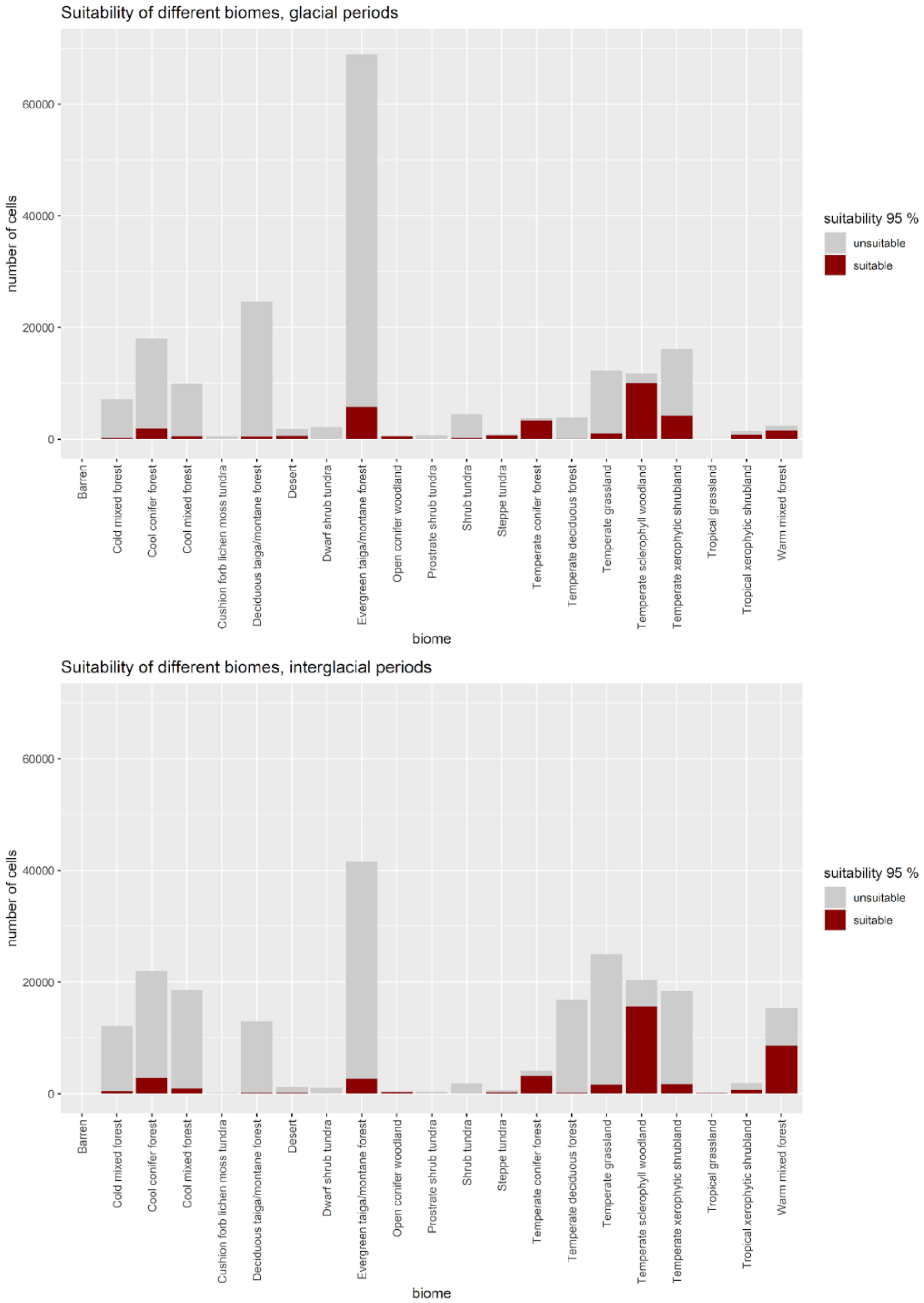


**Supplementary Figure 4**: Amount of habitat suitable for the Acheulean (core and peripheral areas) for each major biome across European forests and shrublands during interglacial and glacial periods (respectively, top and bottom rows).

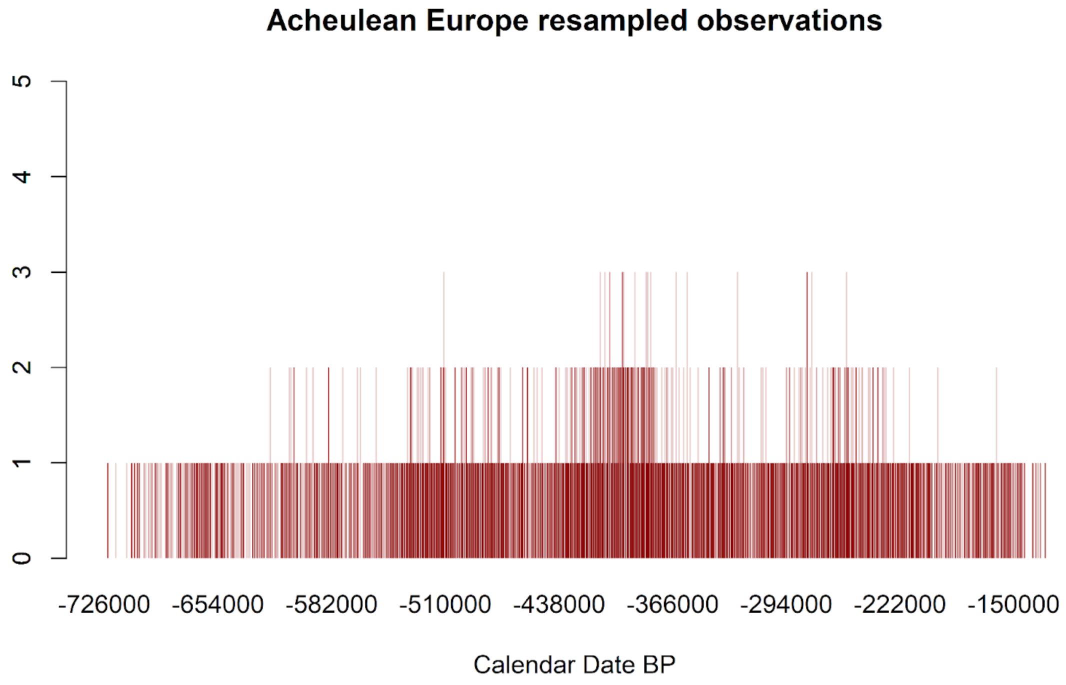


**Supplementary Figure 5**: Chronologically resampled observations. To account for the chronological uncertainty associated with each occurrence, 100 different repetitions of the model were run by sampling a different date from each site's dating range following a normal distribution (mean ± 2-sigma) Each vertical line represents the number of observations in a given time slice and a given repetition. The darker the color, the more repetitions have been sampled from the given time slice.

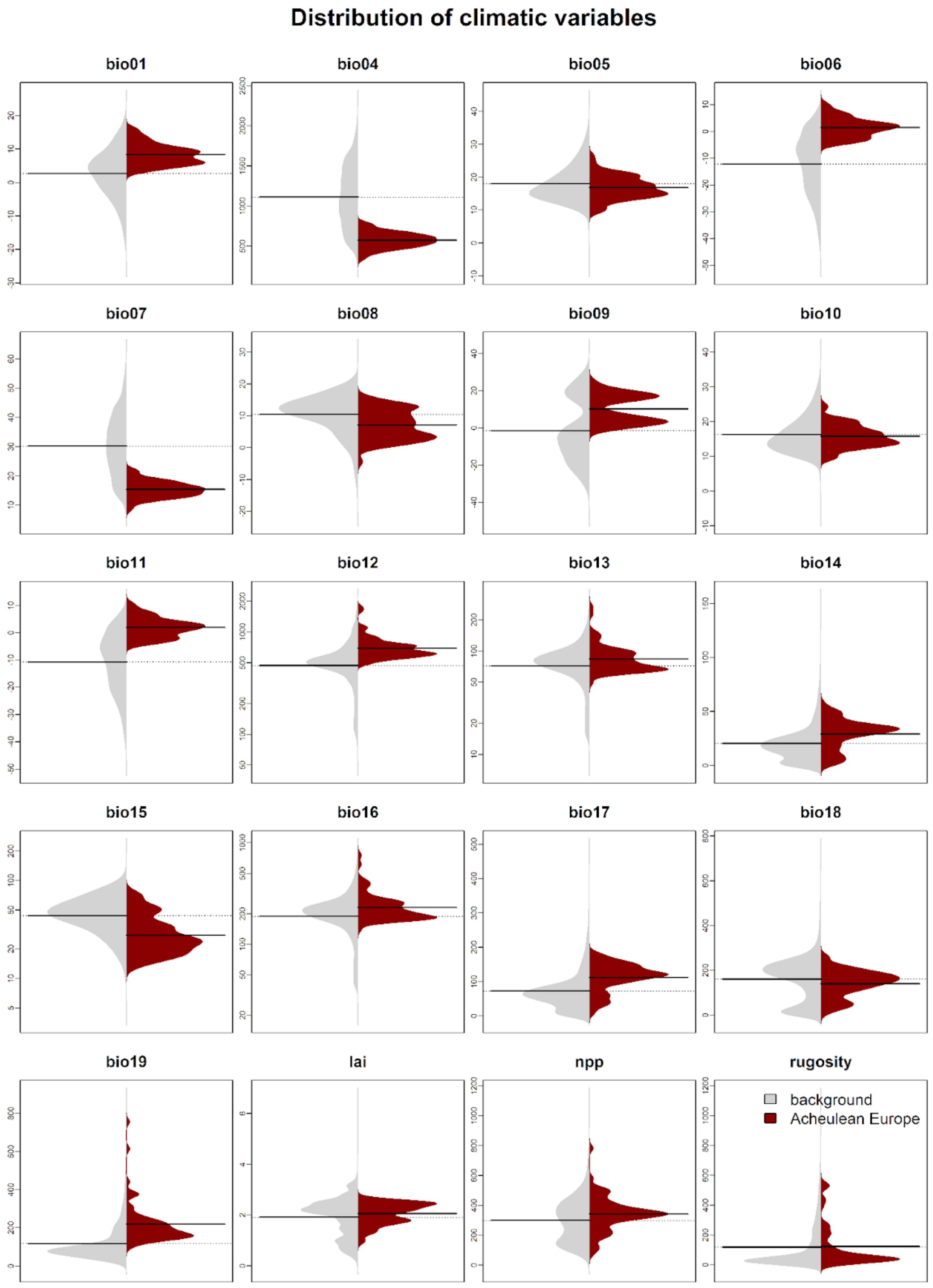


**Supplementary Figure 6:** Violin plots comparing the distribution of each bioclimatic variable in Europe during the whole period considered (background, left side, in light grey) with the one observed at locations where Acheulean sites have been found, considering their mean age (right side, in re
