## Supplementary figures and images for "The Acheulean niche: Climate and ecology predict handaxe production in Europe"

### Supplementary file 1

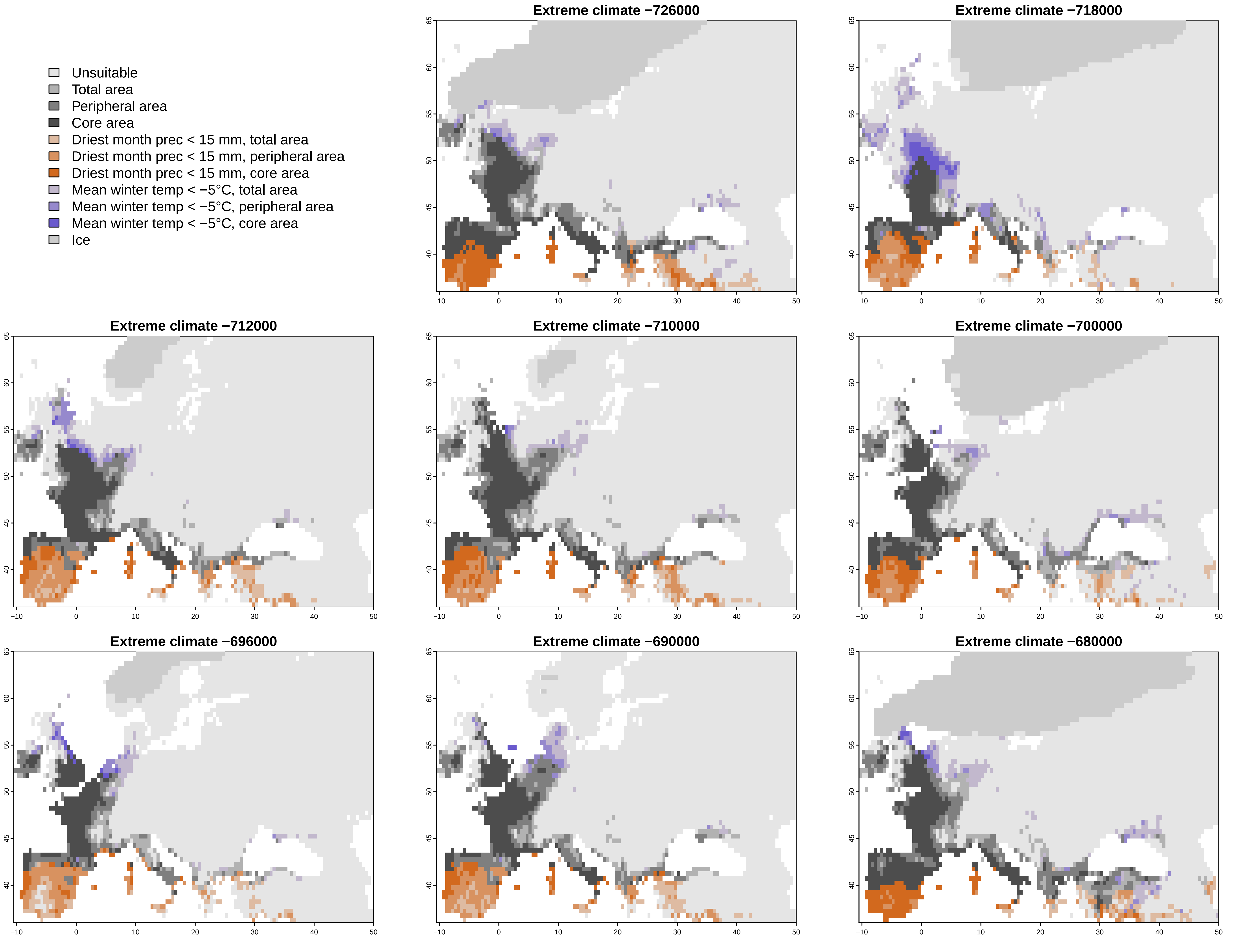

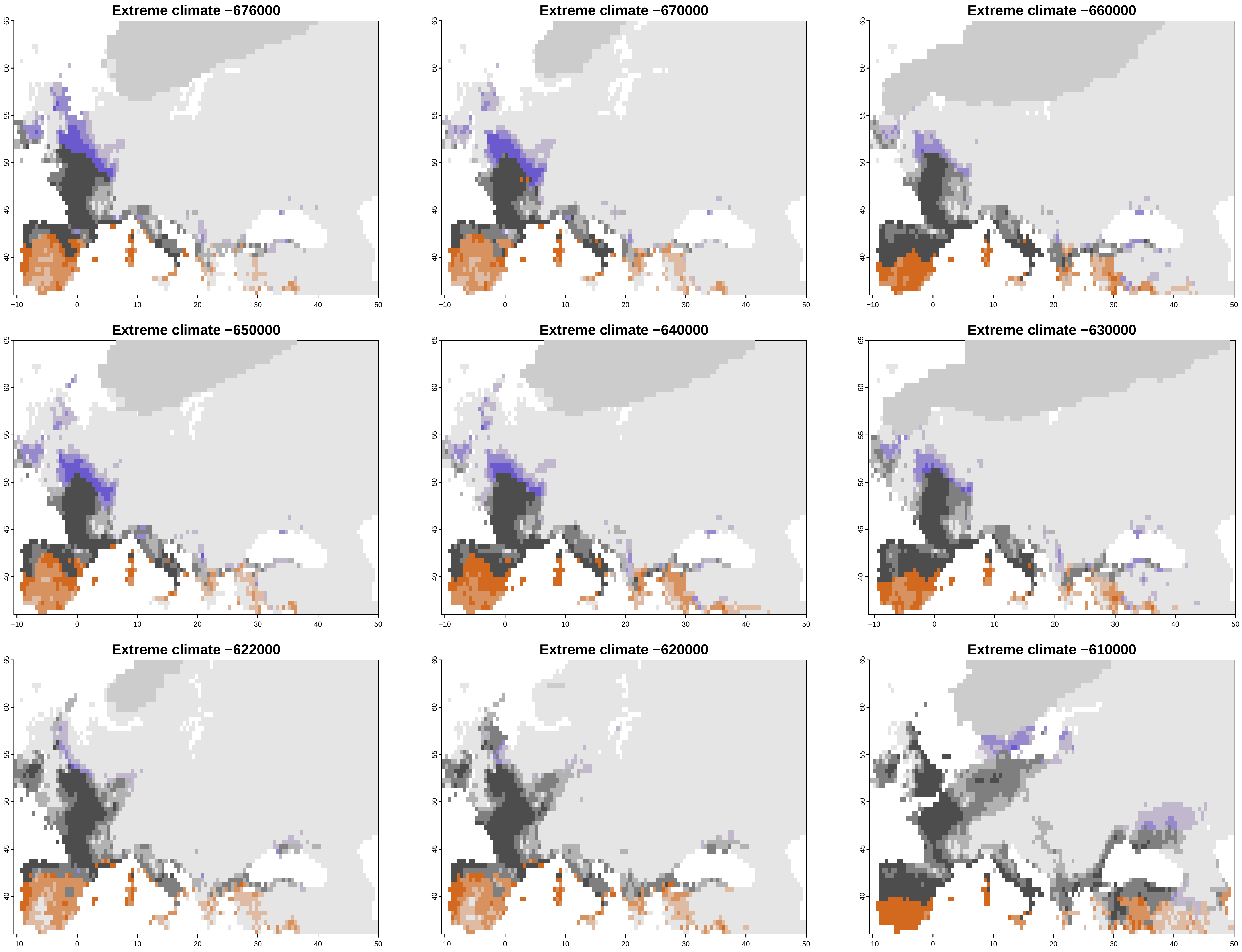

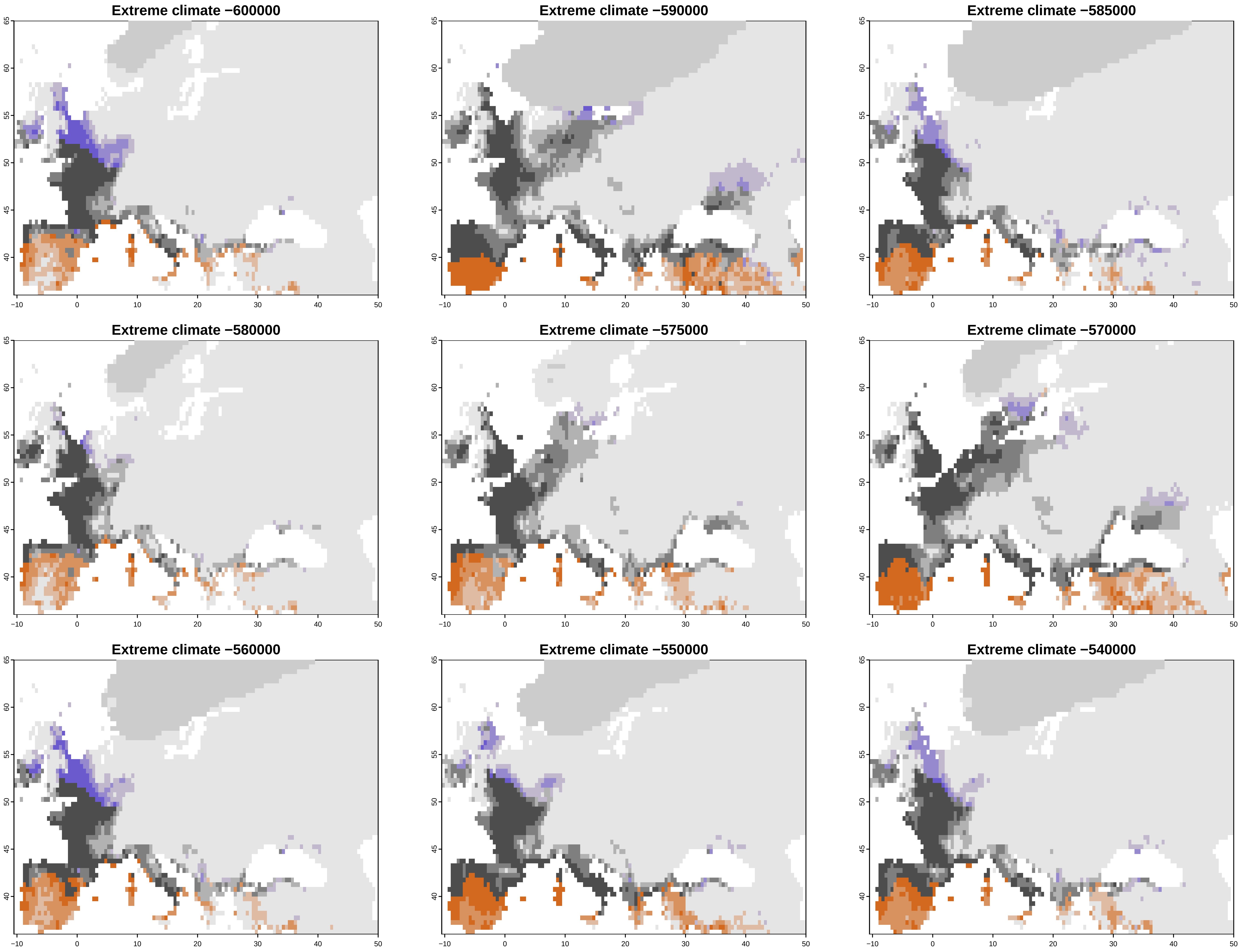

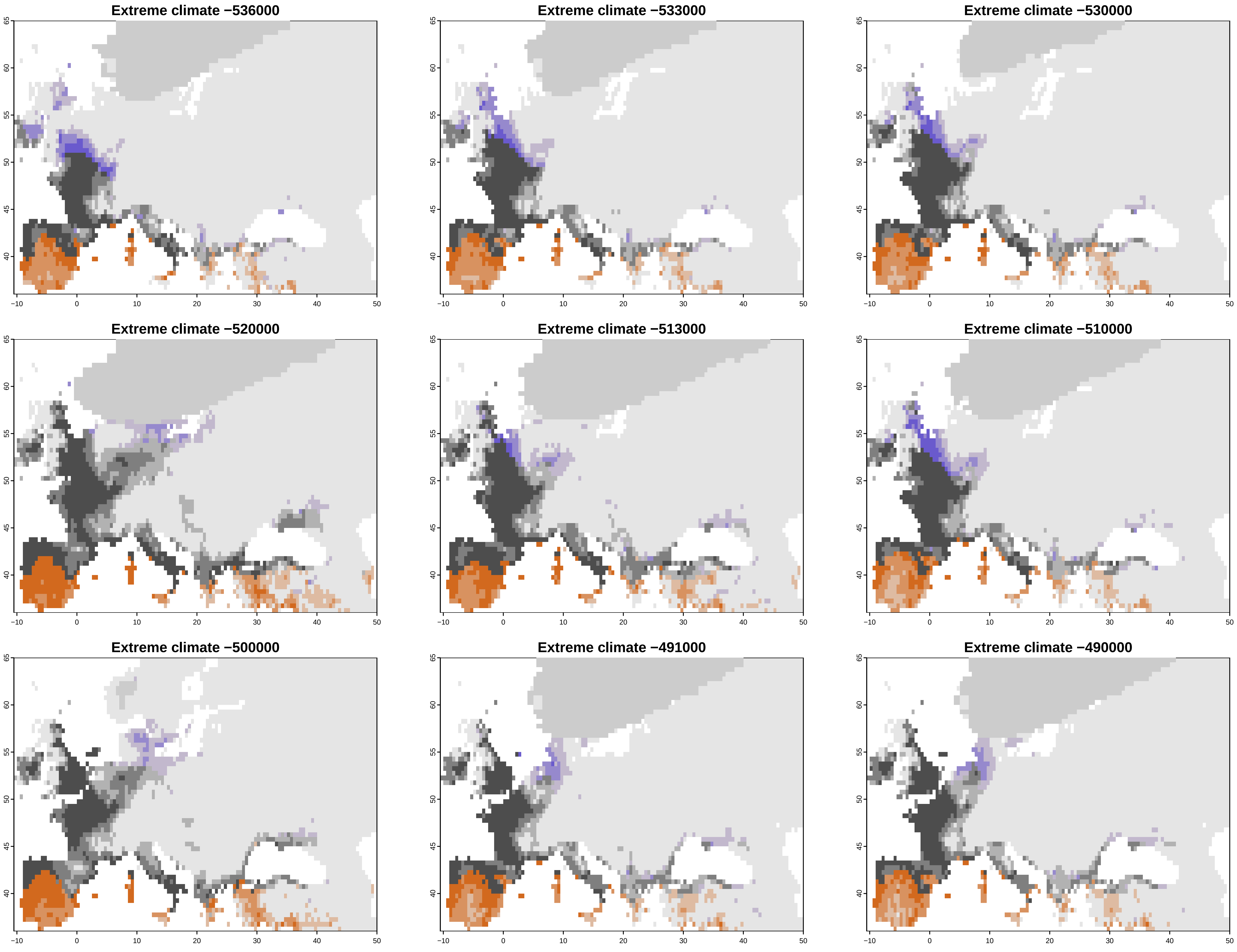

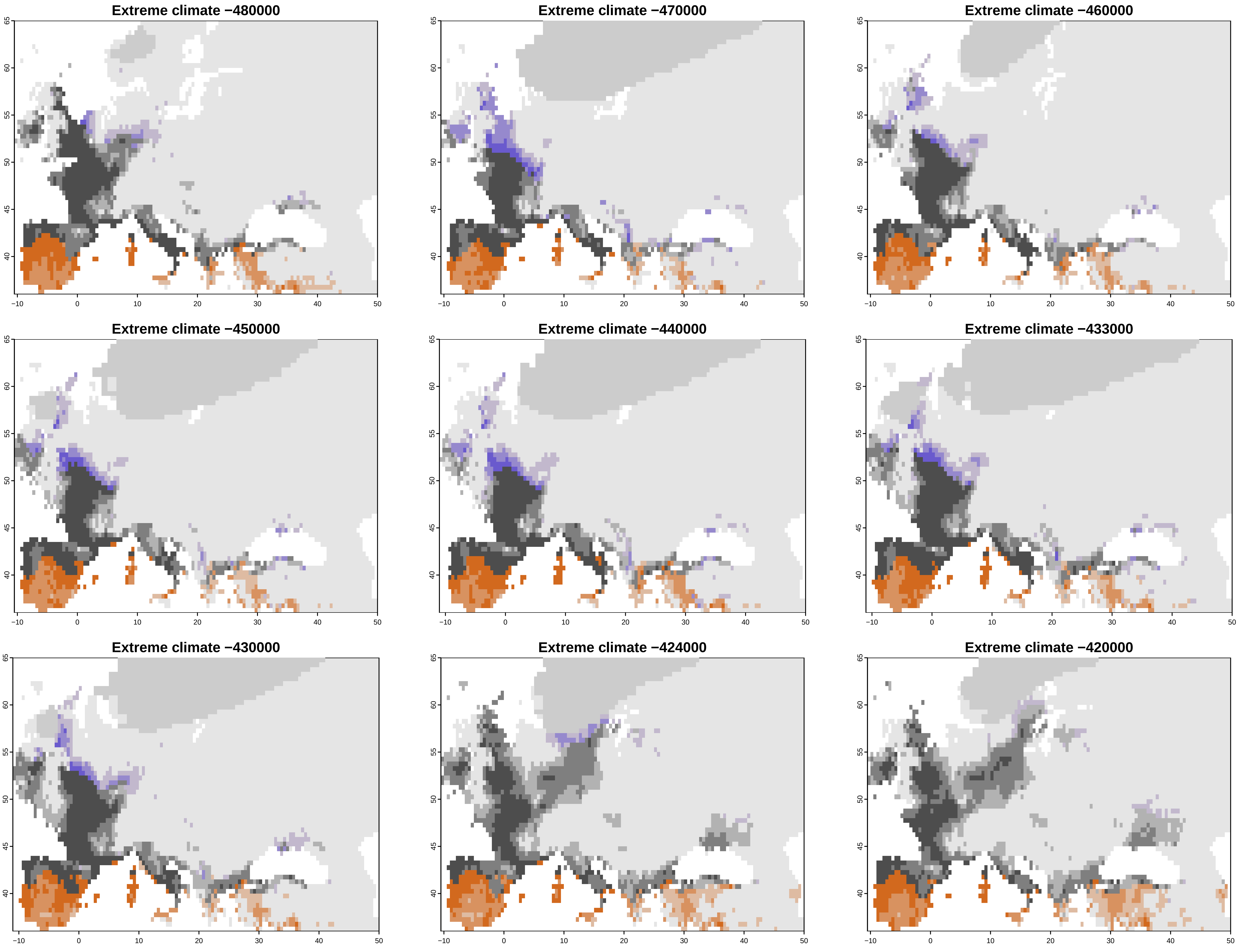

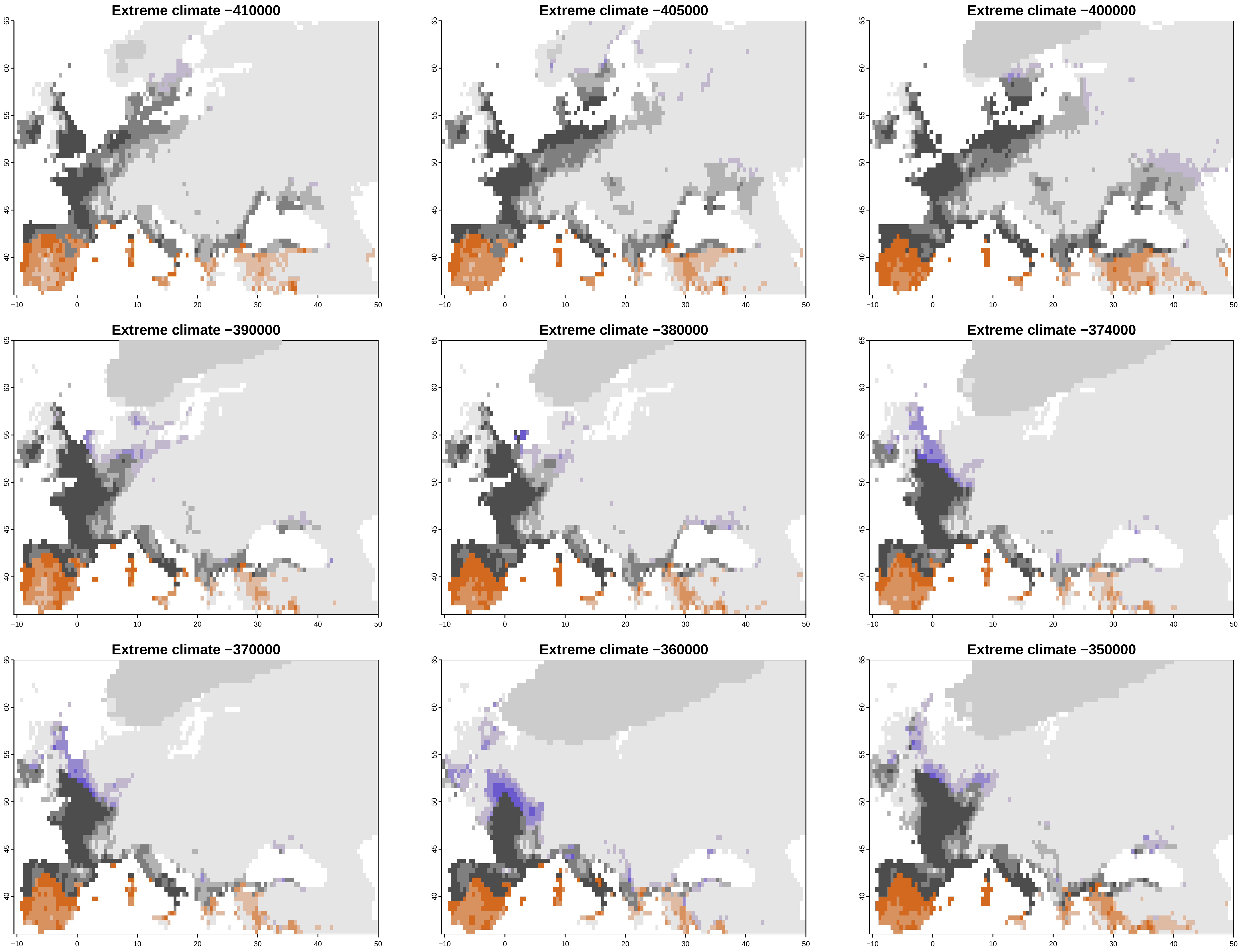

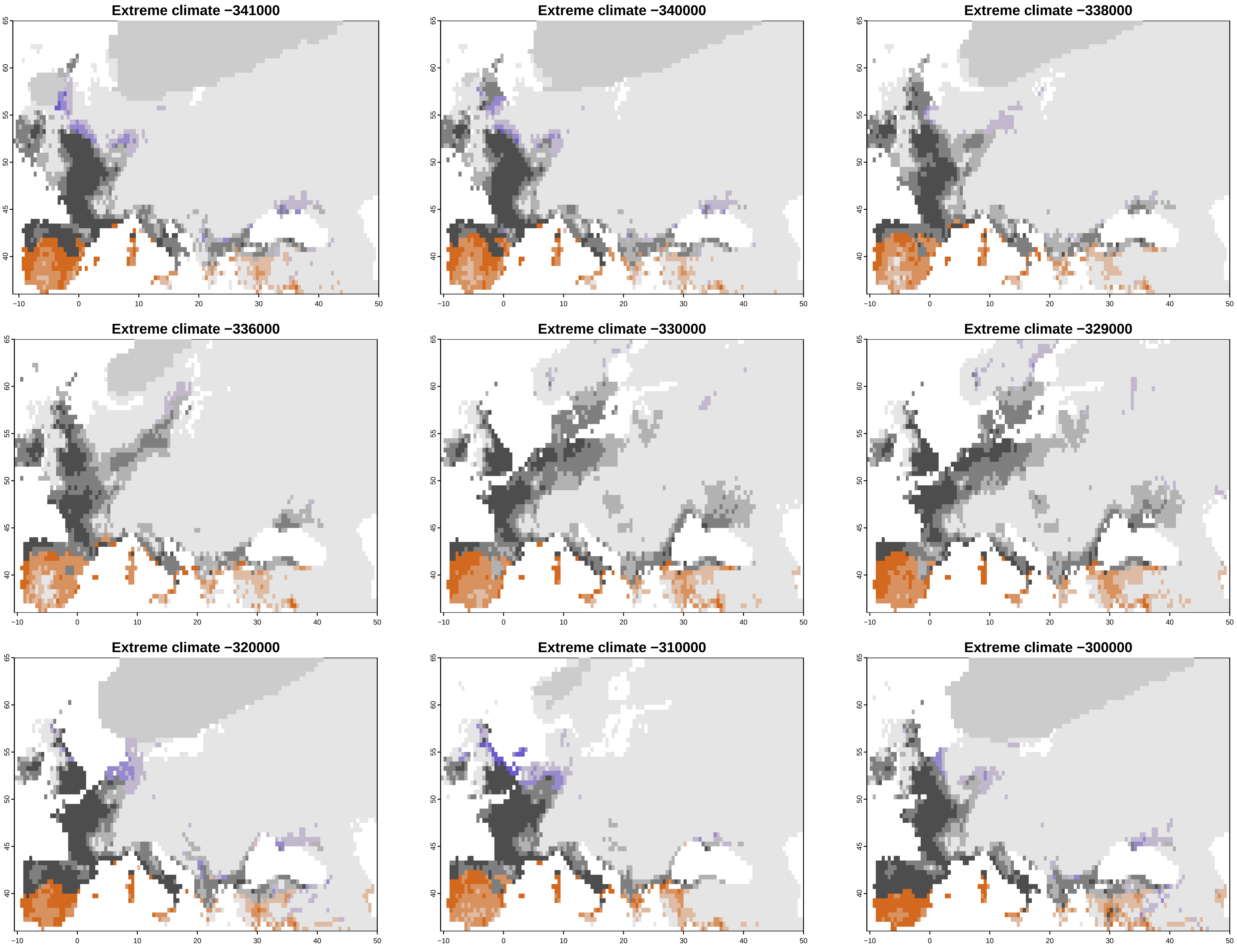

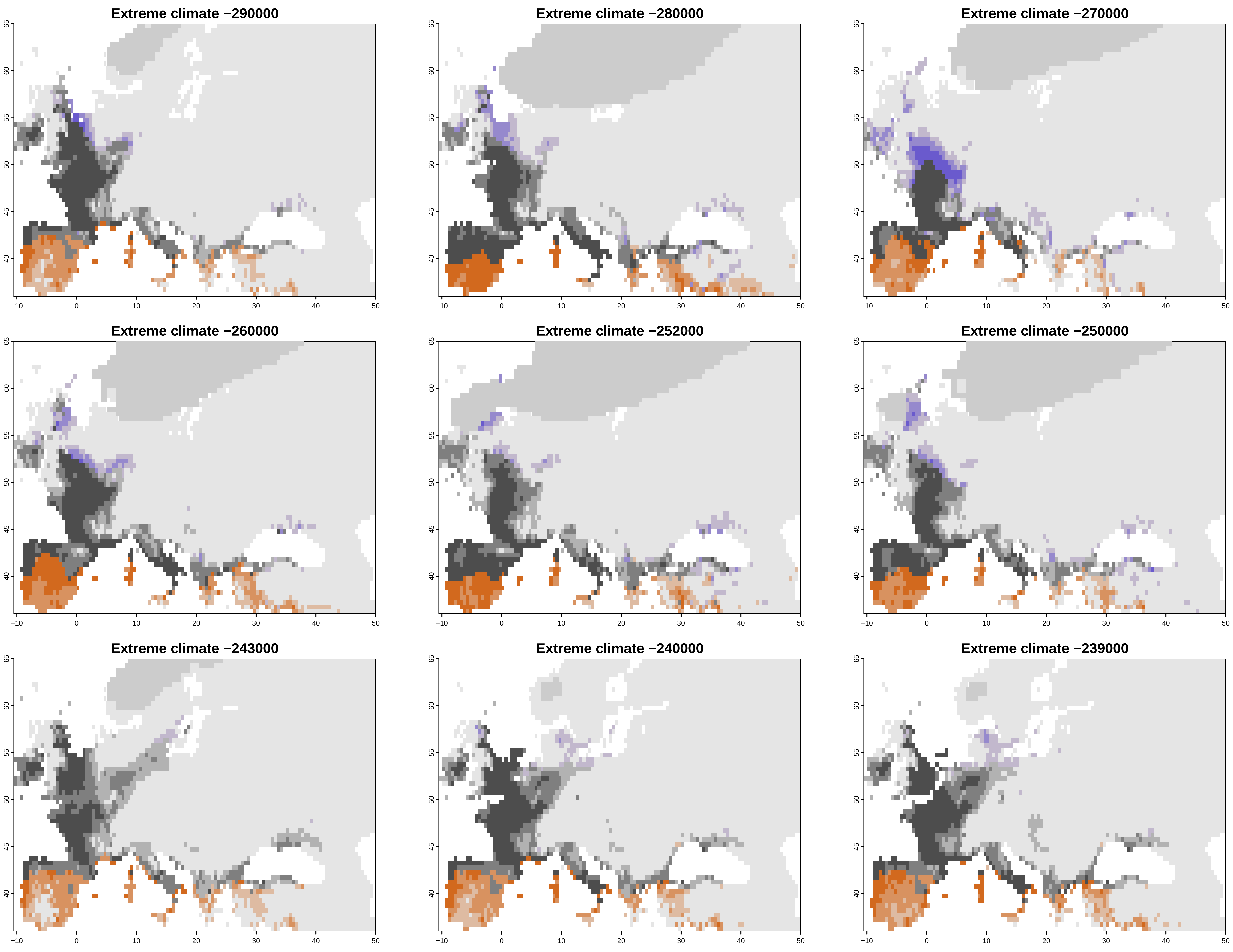

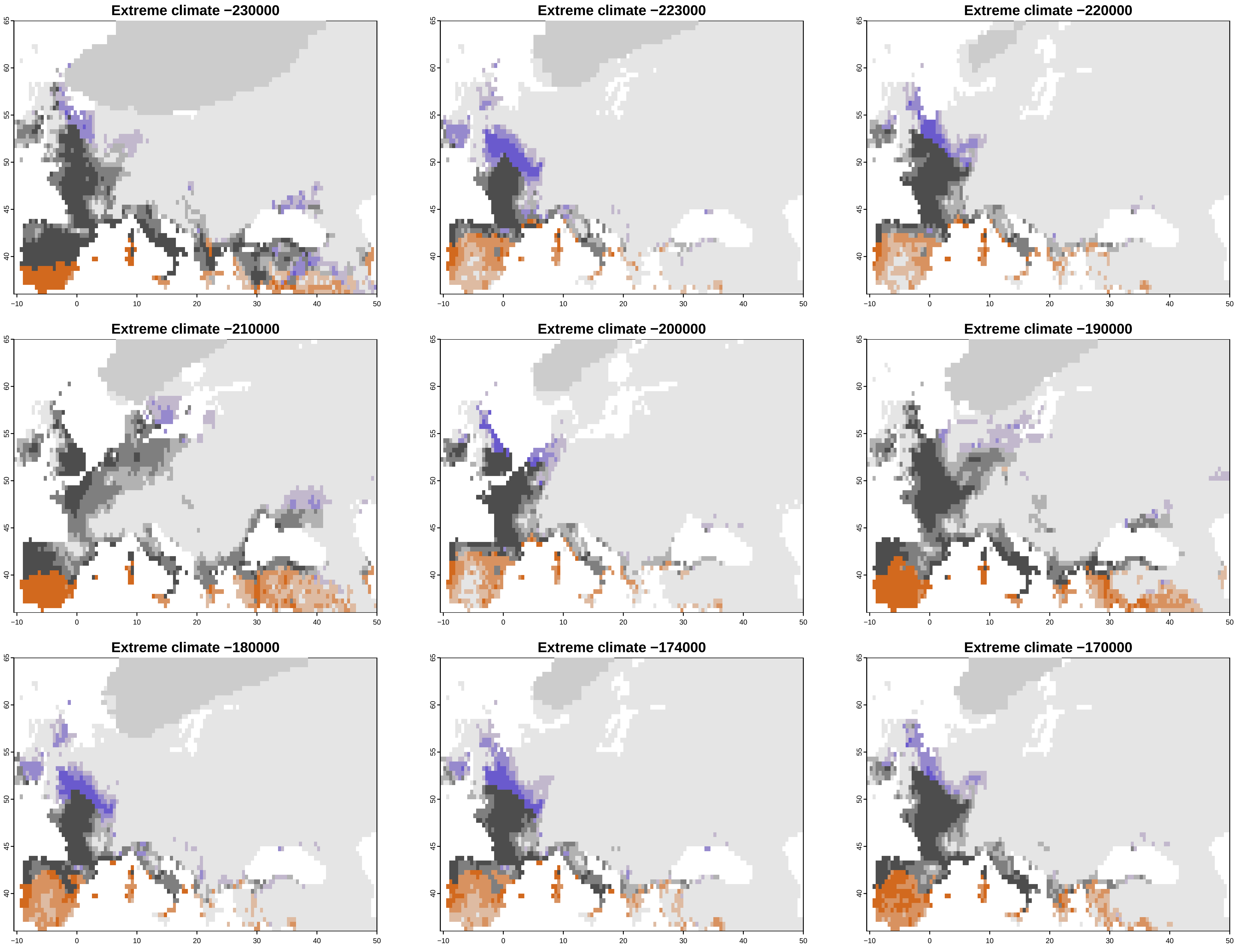

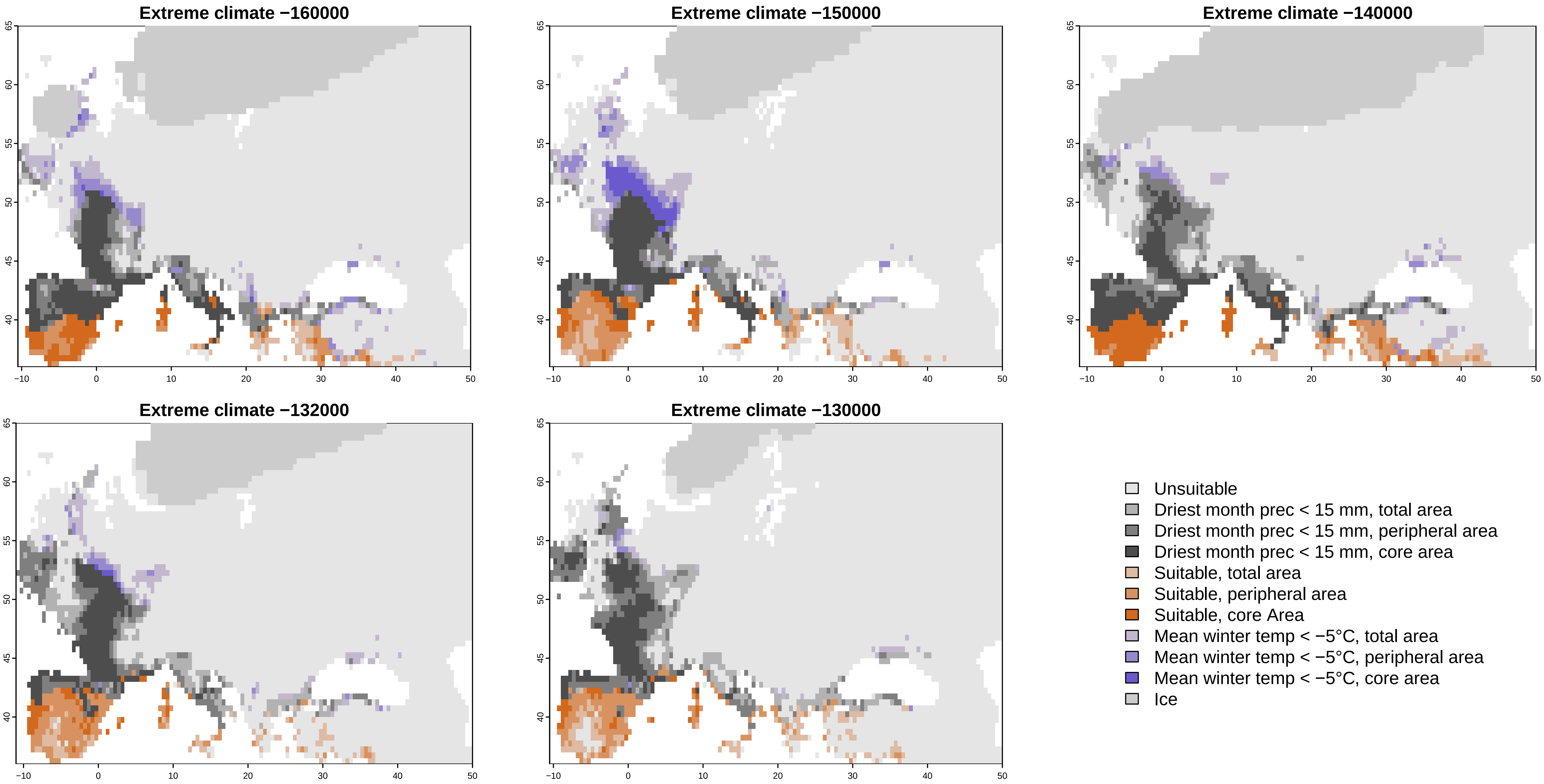

### Supplementary file 2

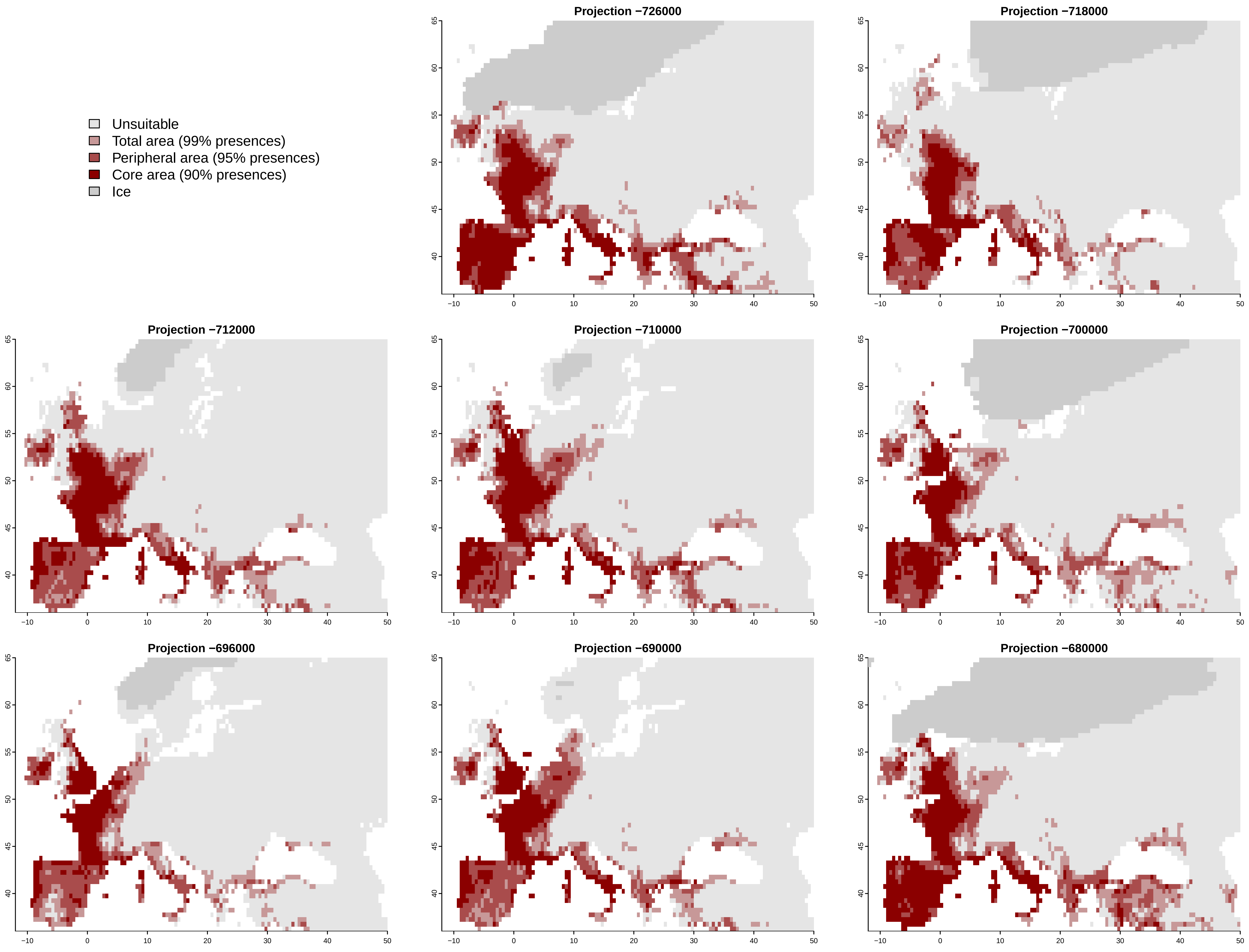

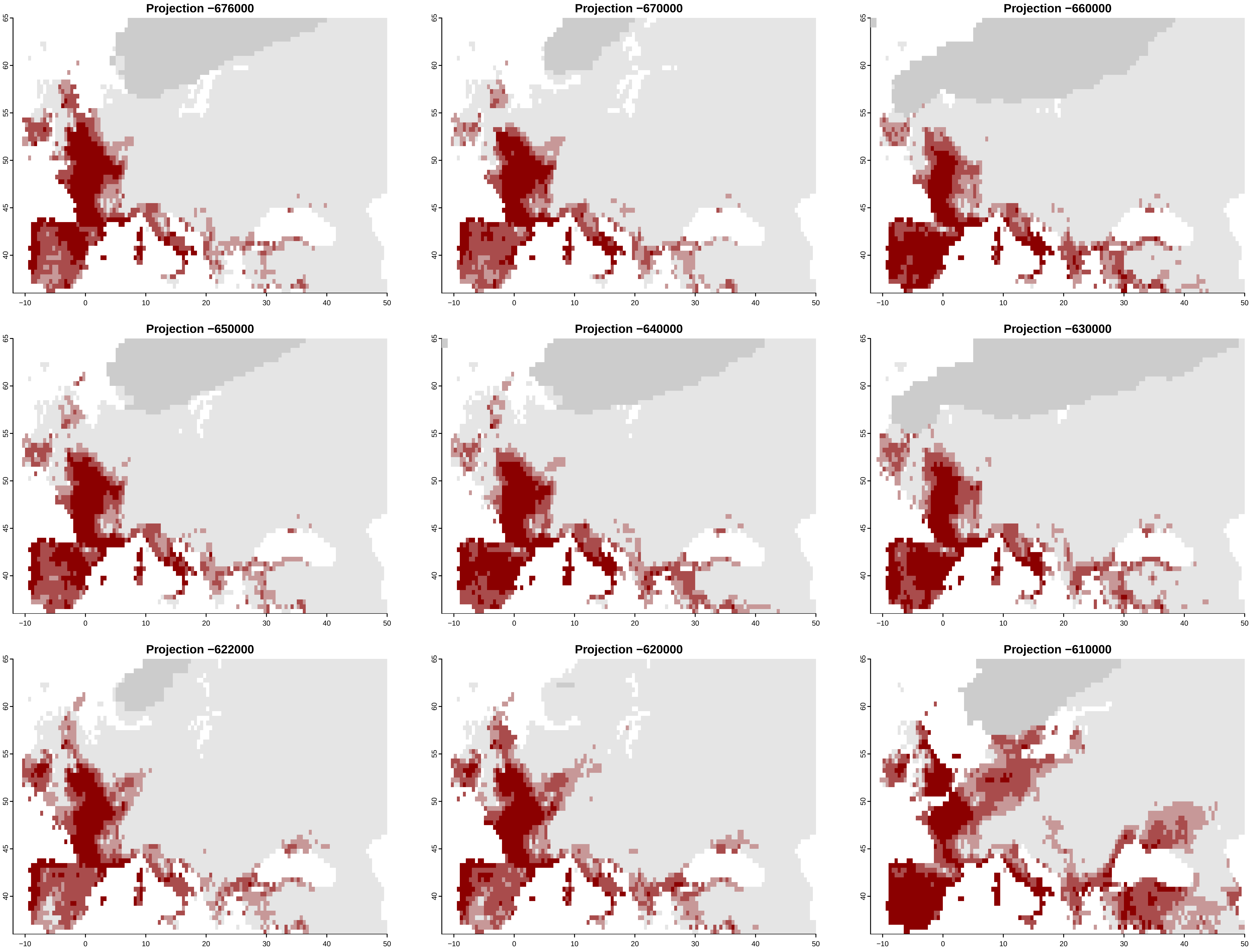

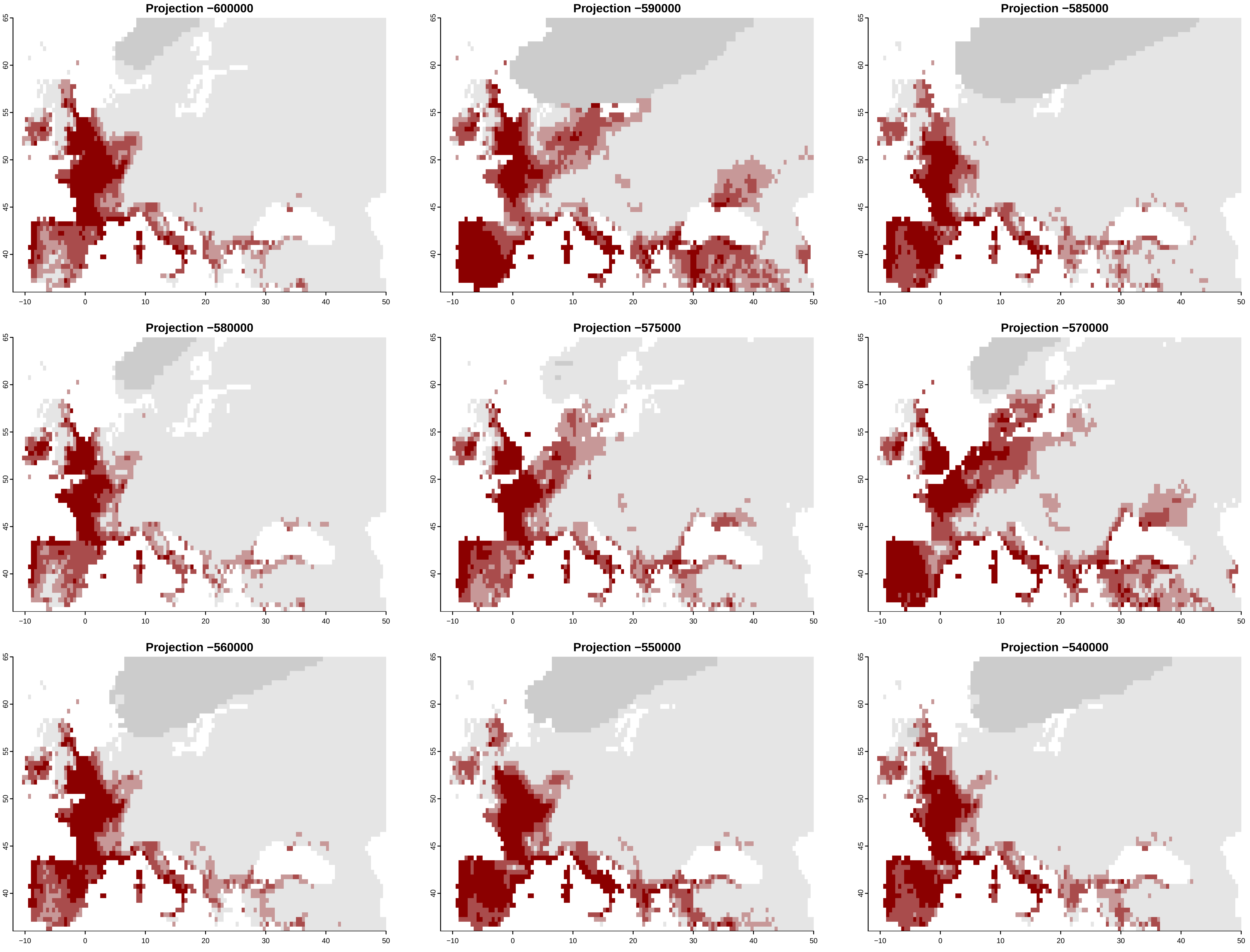

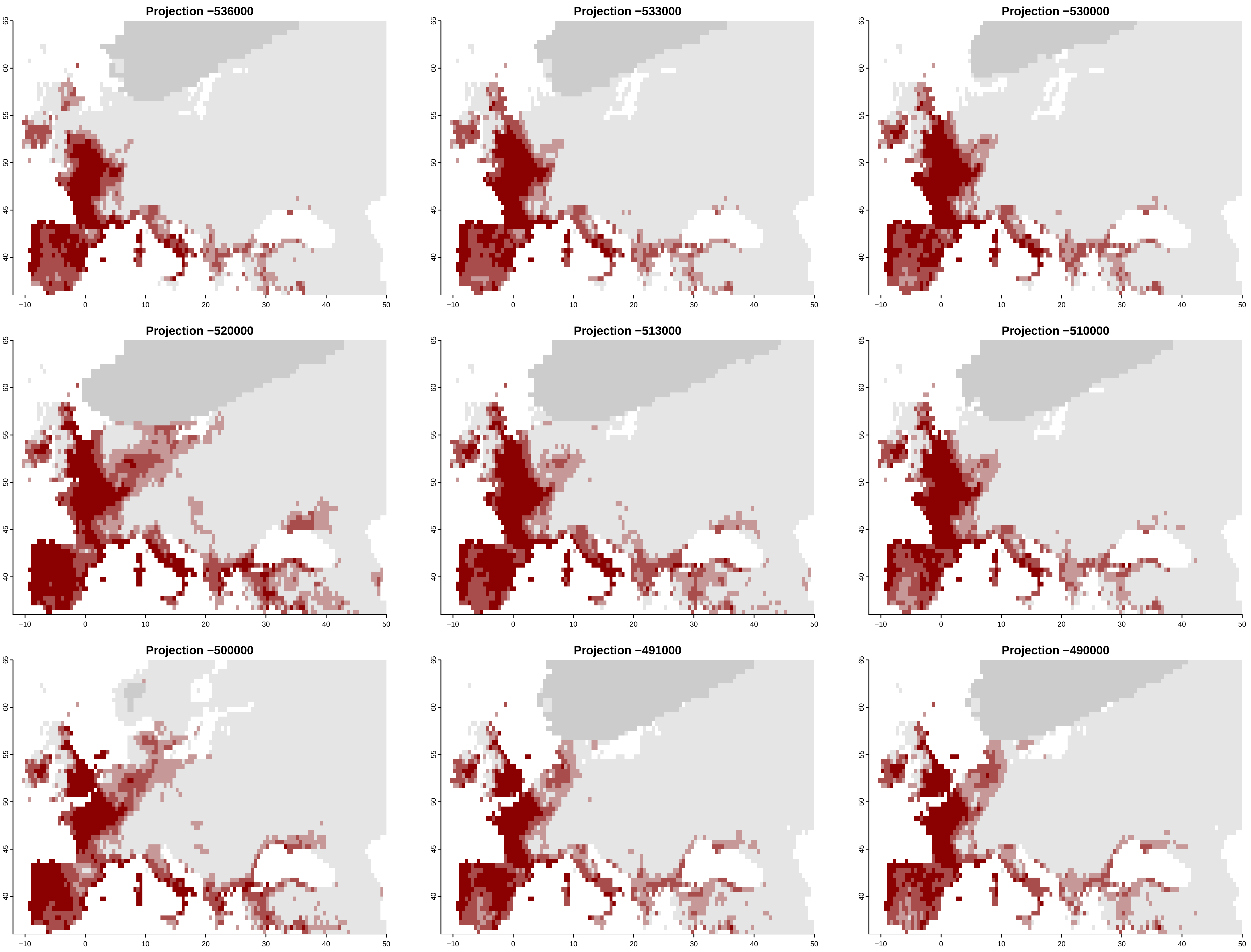

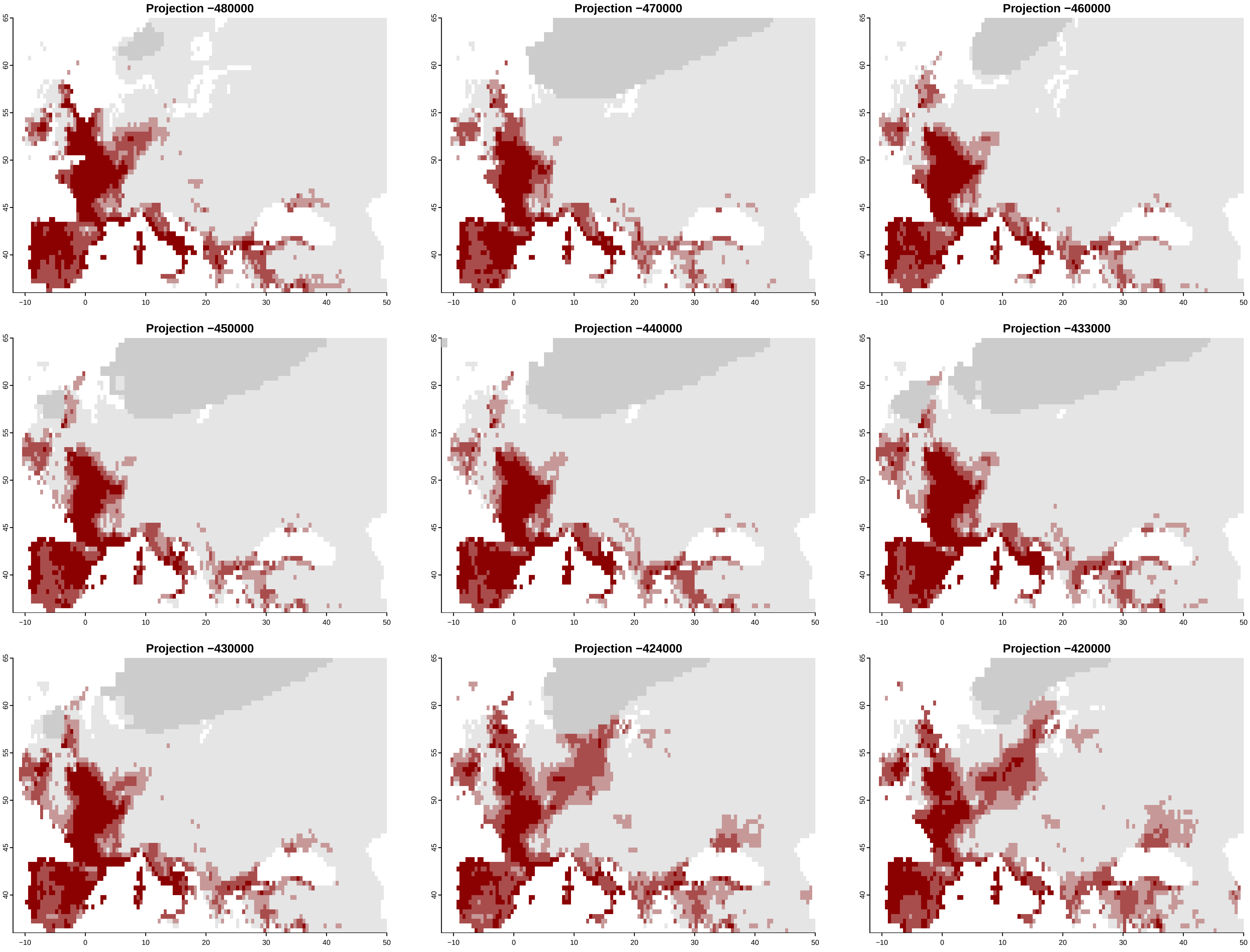

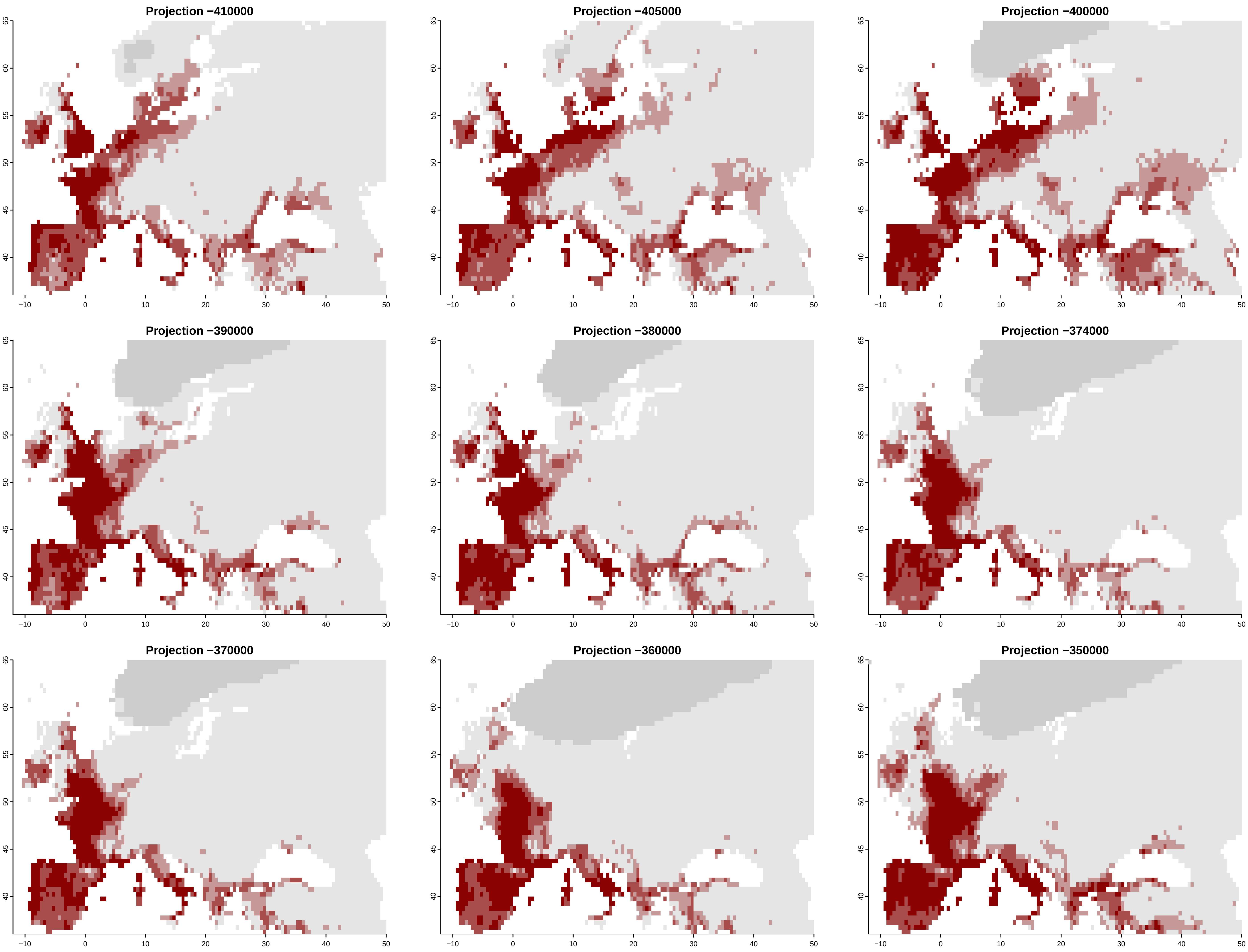

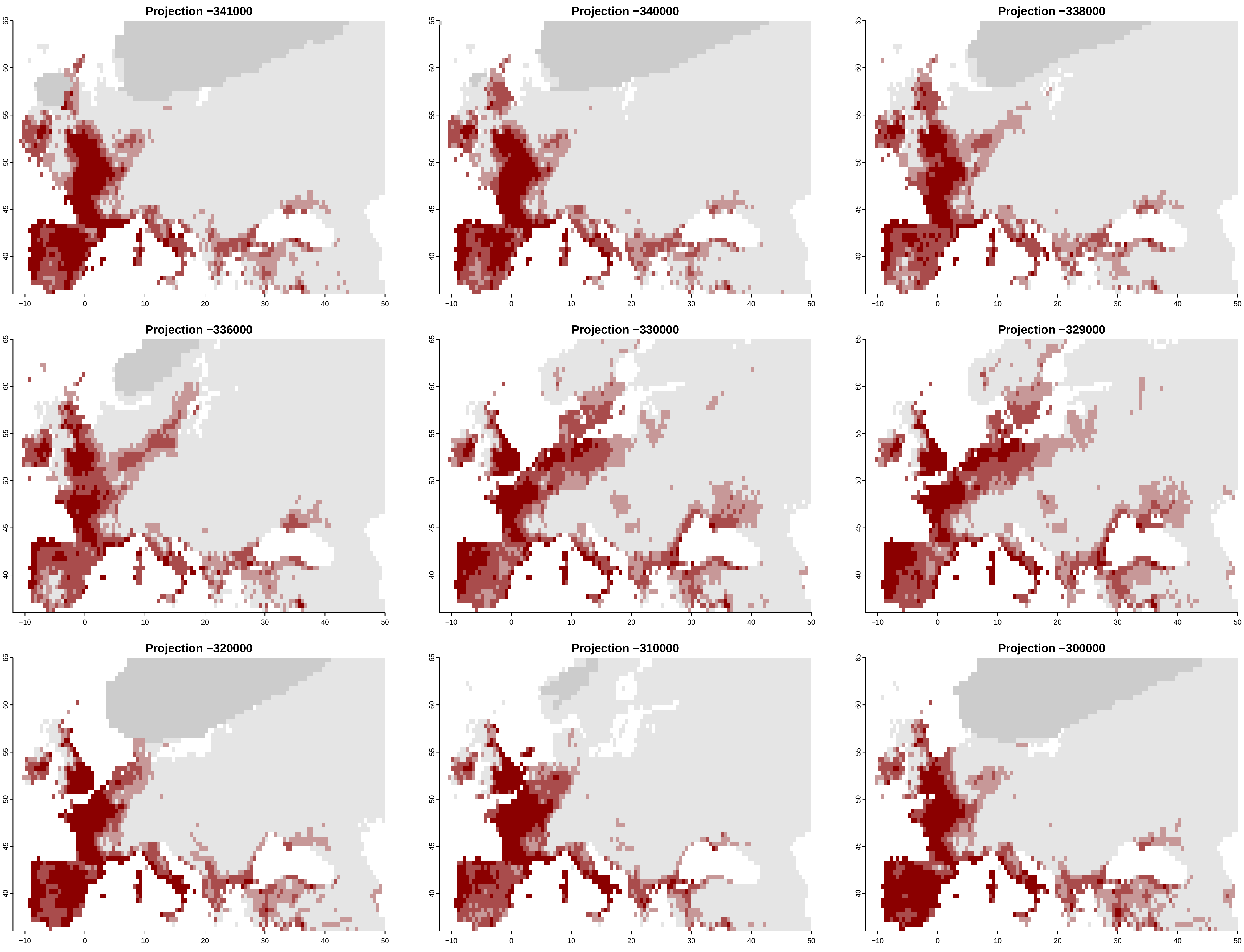

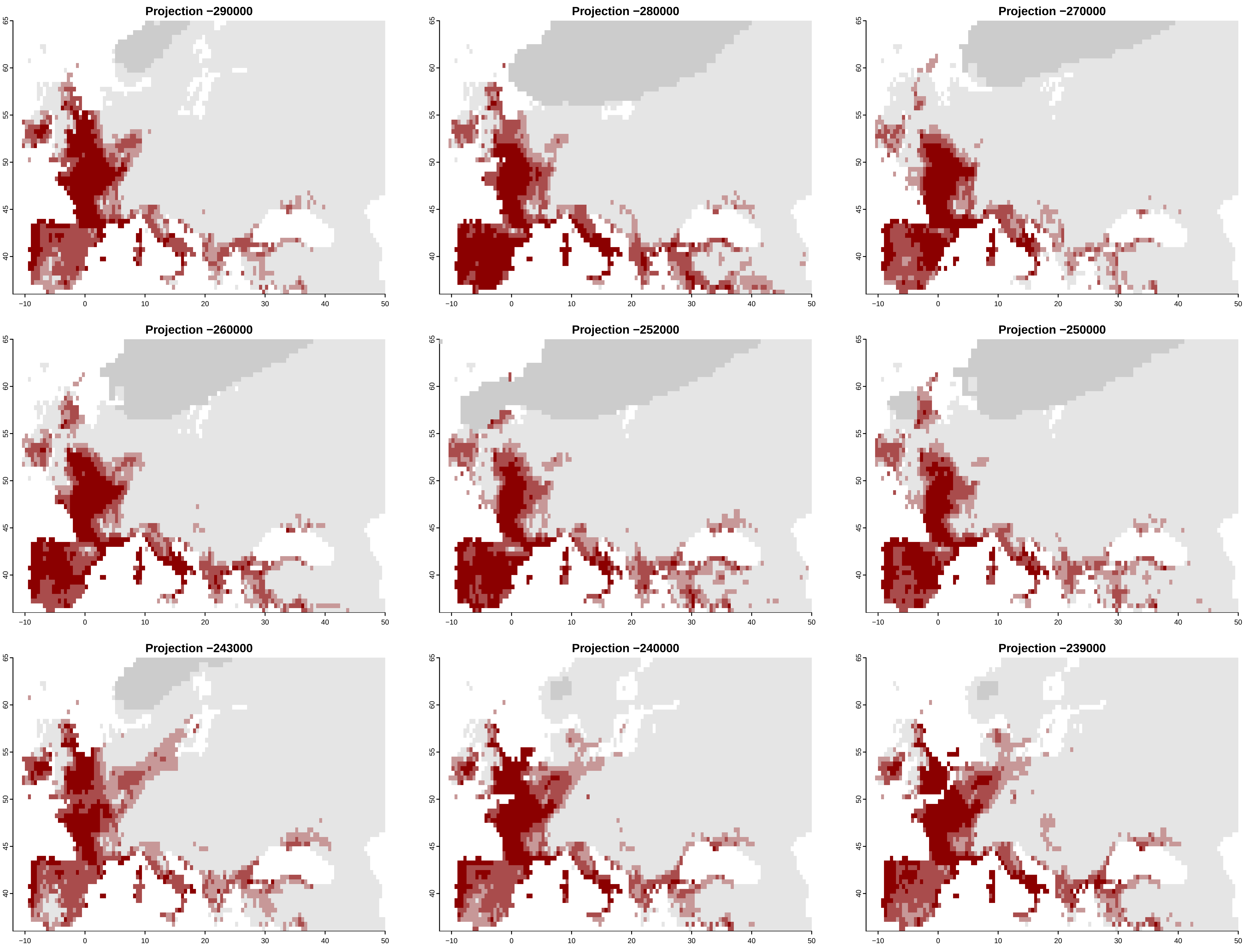

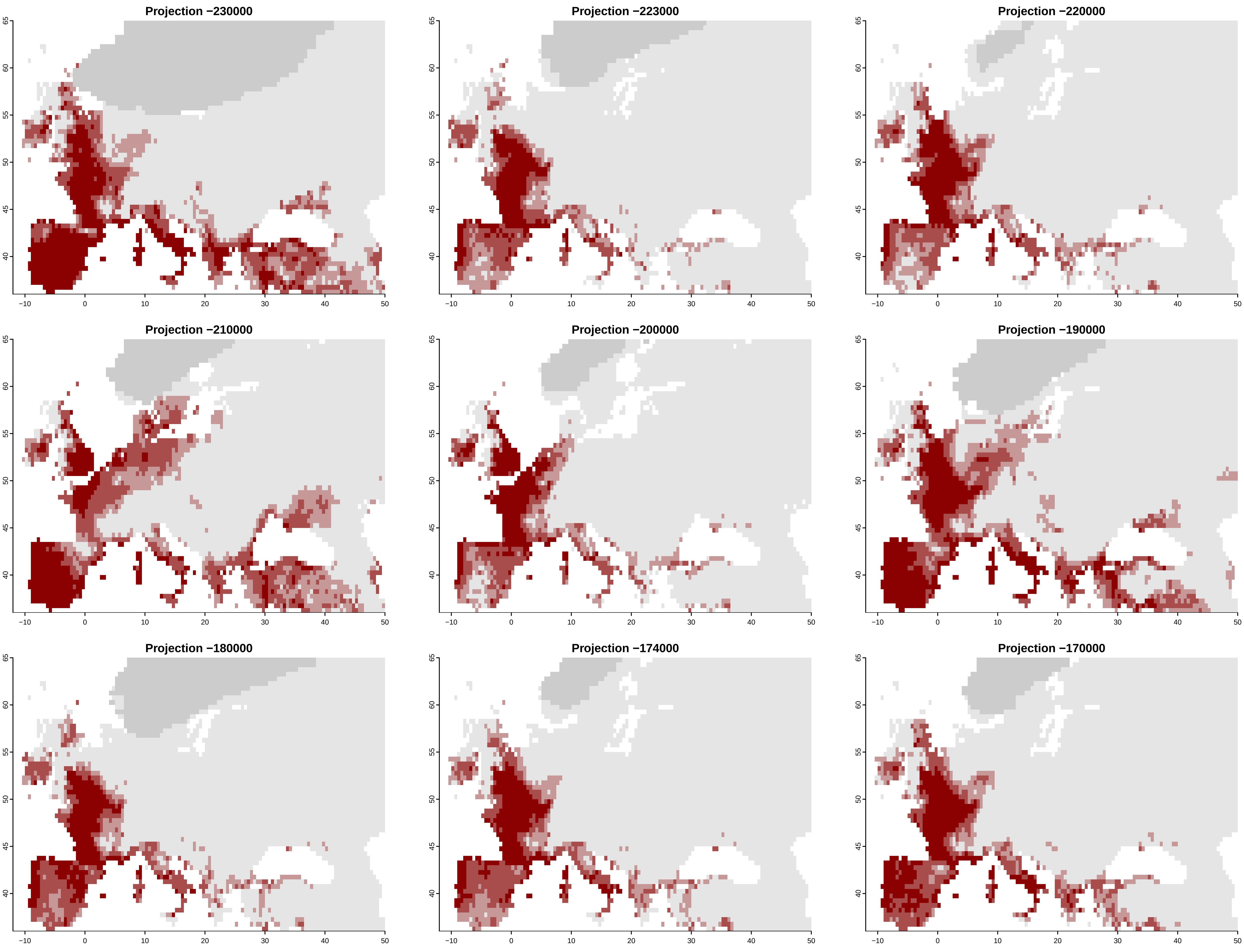

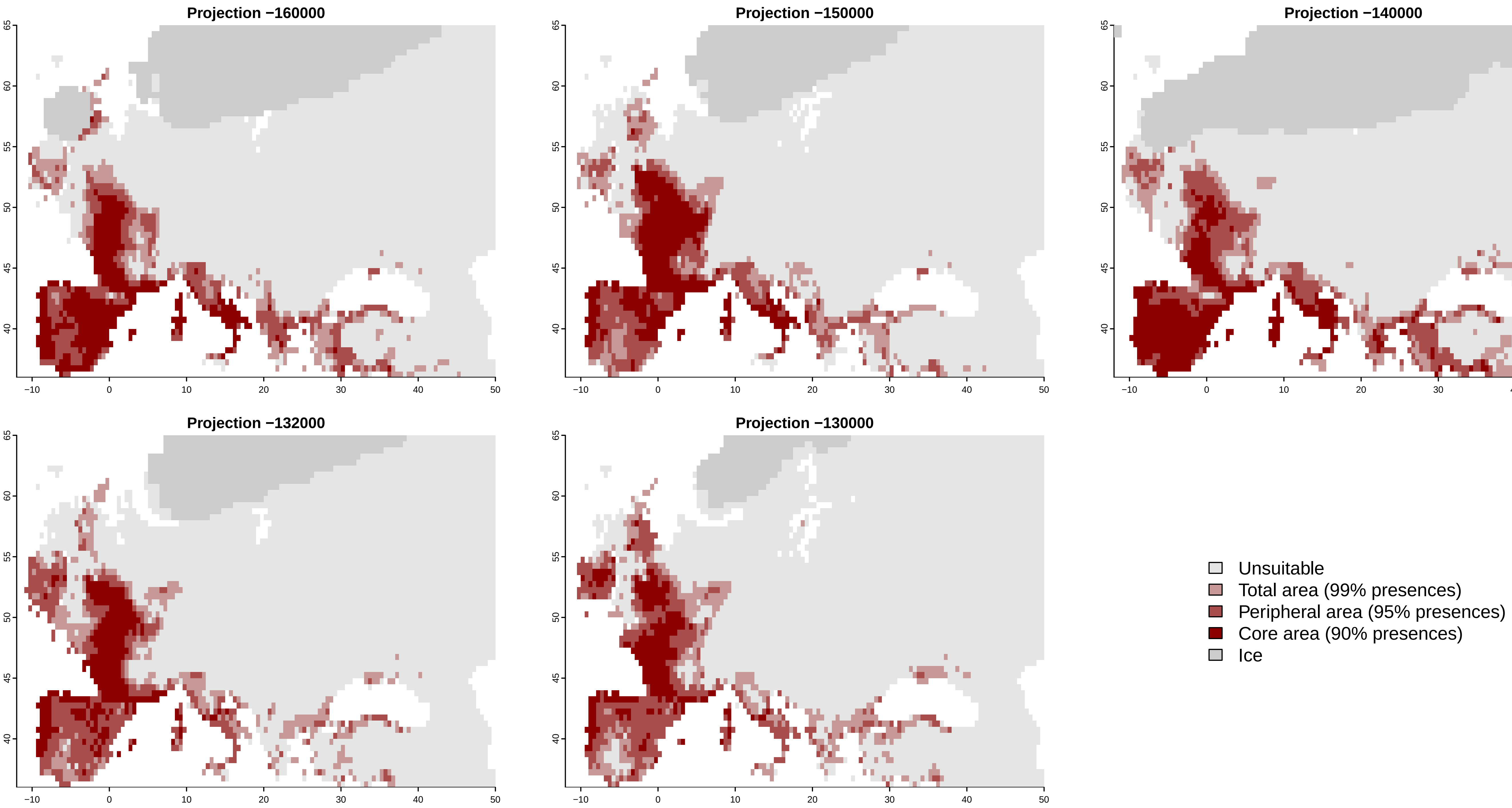
